# Loss of Pigment Epithelium Derived Factor Sensitizes C57BL/6J Mice to Light-Induced Retinal Damage

**DOI:** 10.1101/2024.12.04.626802

**Authors:** Debresha A. Shelton, Jack T. Papania, Tatiana E. Getz, Jana T. Sellers, Preston E. Giradot, Micah A. Chrenek, Hans E. Grossniklaus, Jeffrey H. Boatright, John M. Nickerson

## Abstract

5.1

**Purpose:** Pigment epithelium-derived factor (PEDF) is a neurotrophic glycoprotein secreted by the retinal pigment epithelium (RPE) that supports retinal photoreceptor health. Deficits in PEDF are associated with increased inflammation and retinal degeneration in aging and diabetic retinopathy. We hypothesized that light-induced stress in C57BL/6J mice deficient in PEDF would lead to increased retinal neuronal and RPE defects, impaired expression of neurotrophic factor Insulin-like growth factor 1 (IGF-1), and overactivation of Galectin-3-mediated inflammatory signaling.

**Methods:** C57BL/6J mice expressing the RPE65 M450/M450 allele were crossed to PEDF ^KO/KO^ and wildtype (PEDF ^+/+^) littermates. Mice were exposed to 50,000 lux light for 5 hours to initiate acute damage. Changes in visual function outcomes were tracked via electroretinogram (ERG), confocal scanning laser ophthalmoscopy(cSLO), and spectral domain optical coherence tomography (SD-OCT) on days 3, 5, and 7 post-light exposure. Gene and protein expression of Galectin-3 were measured by digital drop PCR (ddPCR) and western blot. To further investigate the role of galectin-3 on visual outcomes and PEDF expression after damage, we also used a small-molecule inhibitor to reduce its activity.

**Results:** Following light damage, PEDF ^KO/KO^ mice showed more severe retinal thinning, impaired visual function (reduced a-, b-, and c-wave amplitudes), and increased Galectin-3 expressing immune cell infiltration compared to PEDF ^+/+^. PEDF ^KO/KO^ mice had suppressed damage-associated increases in IGF-1 expression. Additionally, baseline Galectin-3 mRNA and protein expression were reduced in PEDF ^KO/KO^ mice compared to PEDF ^+/+^. However, after light damage, Galectin-3 expression decreases in PEDF ^+/+^ mice but increases in PEDF KO/KO mice without reaching PEDF ^+/+^ levels. Galectin-3 inhibition worsens retinal degeneration, reduces PEDF expression in PEDF ^+/+^ mice, and mimics the effects seen in PEDF knockouts.

**Conclusions:** Loss of PEDF alone does not elicit functional defects in C57BL/6J mice. However, under light stress, PEDF deficiency significantly increases severe retinal degeneration, visual deficits, Galectin-3 expression, and suppression of IGF-1 than PEDF ^+/+^. PEDF deficiency reduced baseline expression of Galectin-3, and pharmacological inhibition of Galectin-3 worsens outcomes and suppresses PEDF expression in PEDF ^+/+,^ suggesting a novel co-regulatory relationship between the two proteins in mitigating light-induced retinal damage.

## 5.2 INTRODUCTION

Pigment epithelium-derived factor (PEDF), a secreted 50-kDa glycoprotein with neurotrophic effects, is critical in the development and homeostasis of the vertebrate eye^1–4^. While other ocular tissues express PEDF, the retinal pigment epithelium (RPE) is the primary producer of PEDF and is crucial for retinal health and visual signaling. ^5–9^. RPE ablation studies have shown that loss of the RPE leads to disorganization of multiple retinal layers during development; however, supplementation with PEDF is sufficient to rescue this phenotype in *X. leavis* in ex vivo tissue culture models ^1^. Similarly, loss of the RPE and PEDF expression in the eye is associated with aging^2,10,11^ and ocular pathology^12,13^, including diabetic retinopathy^14,15^ and vascular glaucoma ^15^. PEDF has putative anti-inflammatory roles in eye ^16,17^ and was first described as an anti-tumor factor by Tombran-Tink and colleagues in 1990 because of its ability to differentiate retinoblastoma cells^18,19^. Since then, multiple studies have identified PEDF as a significant support in cellular differentiation, retinal development, inflammation, vascularization, and neuroprotection of photoreceptors and neurons ^7,20–27^. In this study, we asked if PEDF has a protective role in the retina and RPE following LIRD in a C57BL/6J mouse strain that confers resistance to light damage.

In 2006, An et al. studied the secreted proteome of RPE cell cultures isolated from patients with AMD and compared them to control eyes ^28,29^. Interestingly, they found a 3-fold increase in the secretion of four proteins in eye patients with age-related macular degeneration (AMD) compared to controls; among them were galectin-3 (Lgals3) and pigment epithelium-derived factor (PEDF), suggesting that both may be involved in the pathology of the phenotype. Galectin-3, a member of the β-galactosidase binding protein family, is endogenously expressed in the cytosol. Galectin-3 is secreted via a non-classical pathway to the cell surface of the RPE, where it participates in a cell lattice formation and cell-cell interaction observed during EMT of myofibroblastic RPE cells ^3031^ Galectin-3 has also been implicated in fine-tuning inflammatory responses of immune cells during neurodegeneration via its increased affinity for β-1, 6-N-glycosylation on the cell surface of RPE cells undergoing EMT and the increased secretion from RPE and immune cells after damage ^30–35^. However, the role that PEDF expression may play in the modulation of galectin-3 after damage in the eye is not fully understood.

This study identified a novel potential molecular target and signaling pathway that connects the RPE and inflammation via a PEDF-Galectin-3 mediated signaling paradigm. The interplay between PEDF and Galectin-3 may reveal an additional level of regulation of ocular immune privilege facilitated by the RPE over immune cell behavior. Using *in vivo* imaging techniques, electroretinograms, protein and gene expression analysis, and immunofluorescence, we examine how the loss of PEDF expression after light damage increases galectin-3 expression, recruitment of subretinal immune cells, and progressive loss of visual structures and function over time. These findings support the importance of PEDF in protecting eye tissues against LIRD.

## 5.3 METHODS

### 5.3.4 Animal husbandry

The Emory University Institutional Animal Care and Use Committee approved mouse handling, care, housing, and experimental design. The experiments were compiled with the Association for Research in Vision and Ophthalmology (ARVO) and Accreditation of Laboratory Animal Care (AAALAC) guidelines and doctrine. Mice were housed and maintained on a 12-hour light/dark cycle at 22 °C, with standardized rodent chow (Lab Diet 5001; PMI Nutrition Inc., LLC, Brentwood, MO, USA). Mice had access to water *ad libitum*. The Emory University Division of Animal Resources supervised mouse care and housing. A roughly equal representation of male and female mice was used in all experiments. Animals were euthanized using standardized asphyxiation via CO_2_ gas for 5 min, followed by confirmatory cervical dislocation.

### 5.3.5 Breeding Scheme

PEDF knockout/null (PEDF ^KO/KO^ or PEDF-null) mice, which were gifted from Dr. Hans Grossniklaus and Dr. Sue Crawford at Northwestern University Feinberg School of Medicine (JAX Laboratory Stock No. 030065). This mouse strain has had exons 3-6 of the PEDF gene replaced by an IRES-lacZ cassette systemically. We bred PEDF(ko/+) x PEDF(ko/+) on the RPE65 ^M450/M450^ on C57BL/6J . The breeding scheme resulted in litters that were approximately 25% PEDF ^KO/KO^ (experimental) and 25% PEDF ^+/+^ (wildtype controls). These mice were used for all protein and gene expression analysis. To assess immune cell dynamics we used CX3CR-1 ^GFP^ knock-in mice on the C57BL/6J background were acquired from Jackson Laboratory (Stock NO. 005582). We maintained a line that was homozygous for PEDF-ko and another line that was homozygous for PEDF-wt. Both sets of mice were then bred to produce heterozygous CX3CR1(gfp/+) on the RPE65 ^M450/M450^ background. The resultant animals were either PEDF ^KO/KO^; CX3CR-1 ^GFP/+;^ RPE65 ^M450/M450^ or PEDF ^+/+^; CX3CR1 G^FP/GFP^; RPE65 ^M450/M450^. All PEDF ^KO/KO^ experiments were conducted in animals that were more than P60 but less than P380. Genotyping was performed using a polymerase chain reaction to confirm the deletion of the PEDF gene product. The genotyping results were hidden from experimental biologists until after in vivo experiments, and samples were collected to limit ascertainment biases.

### 5.3.6 Light-induced retinal damage (LIRD) conditions and LIRD box information

Mice were dark-adapted overnight before light damage initiation. Phototoxic light damage was induced using Fancier 500-A LED light lamp panels (Fancier Studio, Haywood, CA), which was modified to fit on transparent polycarbonate model 750 cages. The protocol is a modification of previously described phototoxic damage models^36,37^. The light intensity was calibrated using a VWR ® Light Meter with outputs (catalog No. 62344-944, Radnor, PA) to 50,000 lux. The mice were treated with topical 1% Atropine eye drops for two rounds of 10 seconds per eye. Mice were exposed to high-intensity light damage for 5 hours during the dark phase of the animals (7PM-12 AM or ZT12-ZT17). After light damage, animals were returned to their home cages for recovery.

### 5.3.7 Immunofluorescence staining and Histology

#### 5.3.7.1 RPE Flat mounts

Immunofluorescence was used to detect galectin-3 positive cells and RPE cells to assess the extent of immune cell recruitment and damage. Samples were dissected using the technique reported by Zhang et al.^38–40^. In brief, after enucleation, the eye is placed into a 4% Paraformaldehyde/PBS mixture to incubate for 30 minutes. The lens was removed, and four flaps were made to flatten the RPE sheet to a conventional slide with an adhered silicon gasket (Grace Bio-Labs, Bend, OR). The RPE flat mounts were blocked in Hank’s Balanced salt solution (#SH30588.01; Hyclone, Logan, UT) containing 0.3 % (V/V) Triton X-100 and 1% (W/V) bovine serum albumin for 1 hour at 22 °C or overnight at 4°C in a humidity chamber. The samples were then stained with Galectin-3 (1:250), Vimentin (1:250), IGF-1(1:250), and ZO-1(1:200) overnight at 4°C. The next day, the flat mounts were washed with HBSS/Triton X-100 solution and incubated in secondary antibody in HBSS/ Triton 100 X/BSA solution for 1 hour at 22°C. After secondary incubation, samples were washed with HBSS/Triton 100 X solution before mounting with fluoromount G.

#### 5.3.7.2 Retinal Sections

Eyes were fixed in fixation solution (97% methanol, VWR, Cat. #BDH20291GLP; 3% acetic acid, Cat. #Fisher BP2401-500) at −80 °C for 4 days, embedded in paraffin, and sectioned through the sagittal plane on a microtome at thickness of 5 µm as previously described by Sun et al^41^. Nuclei in the outer nuclear layer (ONL) were counted manually by an individual masked to sample identity. Only nuclei within a 100-micron region were counted using Adobe Photoshop (Version 27.4.0) at regularly spaced intervals of 500 microns apart from the optic nerve in both the inferior and superior directions. Deparaffinized retinal sections were also stained for immunofluorescence in a humidity chamber as described by Zhang et al^38^. Slides were mounted using Vectashield Vibrance (Vector Labs H-1700-2; Newark, CA) was used to mount the coverslip, and the sections were imaged using an A1R confocal on a Nikon Ti2 microscope. All primary and secondary antibodies used for this study are listed in Table 1.

**Table 1:** antibody and reagent information.

| Antibody | Antibody Type | Species | Company and Catalog information | Concentration |
| --- | --- | --- | --- | --- |
| **Galectin-3** | Primary | Goat | R&D Systems ( AF1197) | 1:250 |
| **ZO-1** | Primary | Rat | Sigma | 1:250 |
| **Vimentin(D21H3)** | Primary | Rabbit | Cell Signaling (mAB5741S) | 1:200 |
| **IGF-1** | Primary (conjugated AF546) | Mouse | Santa Cruz (sc-518040) | 1:100 |
| **IBA-1** | Primary | Rabbit | Abcam (ab178847) | (1:1000) |
| **Pentahydrate(bis-Benzamide)Hoechst 33258** | DNA nuclear Stain | N/A | Thermo-Fisher Catalog #: H3569 | [1:250] |
| **TUNEL** | N/A | N/A | Promega DeadEnd TUNEL Fluorometric Kit- G3250 |  |
| **Mouse anti-Rhodopsin** | Primary | Mouse | Abcam, ab3267 | [1:250] |
| **Rabbit anti-Best1** | Primary | Rabbit | Abcam, ab14927 | [1:250] |
| **Donkey anti-Rat (AF488)** | Secondary | Rat | Life Technologies, Catalog # A21208 | [1:1000] |
| **Donkey anti-rabbit (AF568)** | Secondary | Rabbit | Life Technologies, Catalog # A10042 | [1:1000] |
| **Donkey Anti-Mouse(AF488)** | Secondary | Mouse | Life Technologies Catalog #A21202 | [1:1000] |
| **Donkey Anti-Goat(AF647)** | Secondary | Goat | Abcam Catalog # A32849 | [1:1000] |

**Table 2:**
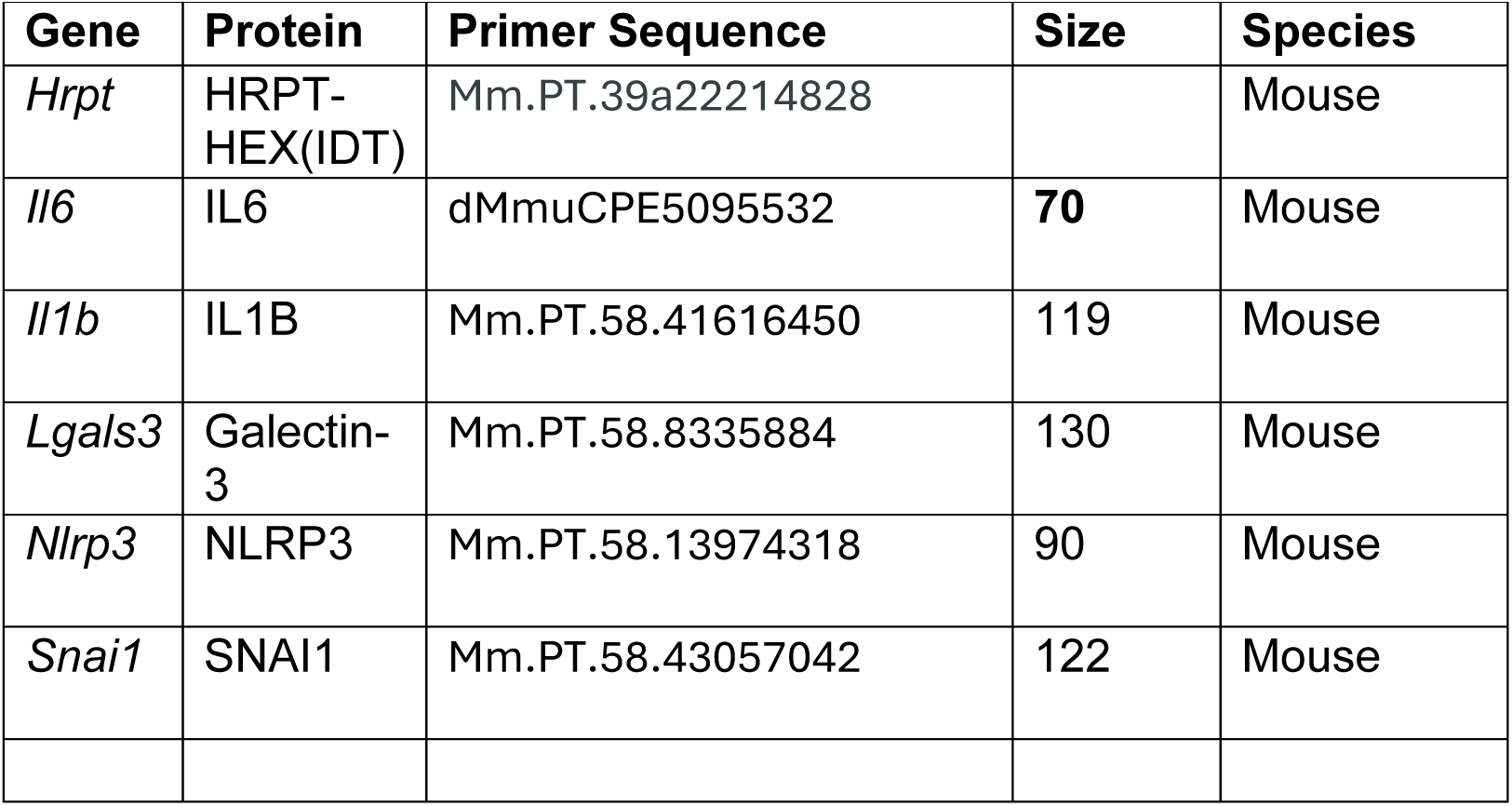
Digital Drop PCR Primer sequences.

| **Gene** | **Protein** | **Primer Sequence** | **Size** | **Species** |
| --- | --- | --- | --- | --- |
| *Hrpt* | HRPT-HEX(IDT) | Mm.PT.39a22214828 |  | Mouse |
| *Il6* | IL6 | dMmuCPE5095532 | **70** | Mouse |
| *Il1b* | IL1B | Mm.PT.58.41616450 | 119 | Mouse |
| *Lgals3* | Galectin-3 | Mm.PT.58.8335884 | 130 | Mouse |
| *Nlrp3* | NLRP3 | Mm.PT.58.13974318 | 90 | Mouse |
| *Snai1* | SNAI1 | Mm.PT.58.43057042 | 122 | Mouse |

### 5.3.8 Rhodopsin staining assay

Animals were euthanized, and eye samples were collected within 1 hour of light onset (between ZT0 and ZT1) to capture maximal phagosome production. Murine eyes were enucleated and placed in glass tubes of “freeze-sub” solution of 97% methanol (Fisher Scientific A433p-4) and 3% acetic acid that was chilled with dry ice, following the method of Sun and coworkers ^42^. Tubes were placed at −80°C for at least four days to dehydrate the tissue. The sections were then treated as described in section 2.4.2. The primary antibodies (mouse anti-rhodopsin, Abcam, catalog #ab3267, [1:250] and Rabbit anti-BEST1, Abcam, catalog # ab14927 [1:250]) are then added to the blocking solution and put on the slides overnight at room temperature in a humidified chamber. The next day, the secondary antibody is added to the blocking solution. Slides were washed and nuclei stained before mounting in fluoromount G (catalog #0100-01; SouthernBiotech, Birmingham, AL, USA). The shed rod outer segments (rhodopsin-positive vesicles) within RPE were quantified as phagosomes. Counts were performed by three independent, masked observers using Photoshop (Adobe Photoshop, Version 27.4.0), and each count was averaged for final counts per sample.

### 5.3.9 Electroretinogram

Mice were dark-adapted overnight for ERG testing, conducted under dim red light conditions as previously described ^43^. Anesthesia was administered intraperitoneally with a 100 mg/kg ketamine and 10 mg/kg xylazine solution ketamine; KetaVed from Boehringer Ingelheim Vetmedica, Inc., Fort Dodge, IA (CAS # 1867-66-9); xylazine from PivetalVet, Greely, CO, USA. Proparacaine (1%; Akorn Inc.) and tropicamide (1%; Akorn Inc.) eyedrops were used for topical anesthesia and pupil dilation. Mice were kept on a 39 °C heating pad during the procedure. ERGs were recorded using the Diagnosys Celeris system (Diagnosys, LLC, Lowell, MA, USA), with corneal electrodes on each eye and the contralateral eye as the reference. Full-field ERGs were recorded for scotopic conditions at stimulus intensities of 0.001, 0.005, 0.01, 0.1, and 1 cd s/m² with a 4 ms flash duration, collecting signals for 0.3 sec to assess a- and b-wave function. For c-wave analysis, a 10 cd s/m² flash was used, with a 5-sec signal collection. After light adaptation for 10 minutes, photopic ERGs were captured at 3 and 10 cd s/m². Post-recording, mice were placed in their home cages on heating pads to recover from anesthesia unless further prepared for SD-OCT and cSLO examinations.

### 5.3.10 In Vivo Ocular Imaging

#### 5.3.10.1 Spectral Domain Optical Coherence Tomography (SD-OCT)

Mice were anesthetized during the previous ERG examination, and a ketamine booster was administered to extend the examination period. The procedure for in vivo ocular posterior segment morphology analysis has been previously described ^38^. In brief, spectral domain optical coherence tomography (SD-OCT) using the MICRON^®^ IV Spectral Domain Optical Coherence Tomography (SD-OCT) system with a fundus camera (Phoenix Research Labs, Pleasanton, CA, USA) was used sequentially to examine the retinal anatomy. Micron IV system, circular scans ∼100 µm from the optic nerve head were collected (50 scans averaged together) to generate image-guided OCT images of retinal layers and fundus. Retinal layers were annotated according to previously published nomenclature ^44^Total retinal thickness and photoreceptor (outer nuclear layer thickness) were analyzed using Photoshop (Adobe Photoshop 2024 version 25.5) as previously described^38^.

#### 5.3.10.2 Confocal Scanning Laser Ophthalmoscope (cSLO)

Immediately afterward, a rigid, specialized contact lens adapted for mouse imaging (Heidelberg Engineering) was placed on the eye (back optic zone radius, 1.7 mm; diameter, 3.2 mm; power, Plano), and blue autofluorescence (BAF) imaging at the layer of the photoreceptor-RPE was obtained using Heidelberg Spectralis and SD-OCT instrument with a 25 D lens (HRA)CT2-MC; Heidalberg Engineering, Heidalberg, Germany). Afterward, mice were injected with a reversal agent (0.5 mg/mL atipamezole(Antisedan); Zoetis, Parsippany, NJ) injection volume 5 µL per gram mouse weight; and placed individually in cages on top of heated water pads to recover.

### 5.3.11 Western Blot Protocol

As described in Ferdous et al. 2019 and Ferdous et al. 2023, immunoblot experiments were conducted. In brief, two dissected eye cups (containing both the retina and RPE/ Sclera) were collected from each animal. Protein was extracted via mechanical rending of tissue by a QIAGEN TissueLyser in a solution of radioimmunoprecipitation (RIPA) buffer containing protease inhibitors (completed mini protein inhibitor catalog #118361530001) and phosphatase inhibitors (PhosSTOP EASypack #04906845001). Protein concentration was determined using Pierce bicinchonic Acid (BCA) Assay, and absorbance was measured at 562 nm using a Synergy H1 Hybrid Plate Reader (Biotek). After ascertaining protein concentration, the samples were diluted to 0.8 mg/mL and heated to 95 °C for 10 minutes to denature proteins before electrophoresis. Samples were run on a pre-cast Criterion gel (Biorad TGX Stain free Gel 4%-20%, Catalog # 567-8094) along with 10µL of a molecular weight ladder (Bio-Rad Catalog # 1610376) and run at 120V for 90 mins.

### 5.3.12 TUNEL Staining protocol

The manufacturer instructions for the Promega DeadEnd TUNEL Fluorometric kit (Promega G3250) were followed. In brief, tissue sections were deparaffinized in 5 steps of xylene for 2 min each. The tissue sections were then rehydrated in a graded ethanol series (100, 90, 80, 70, 60, and 50%) for 2 min each. The slides were then washed for 5 min in PBS (Corning 46-013-CM) and mounted in the Sequenza system. Sections were incubated for 15 min in Z-fix (Anatech, Fisher Scientific NC935141), washed twice in PBS for 5 min each, incubated in Proteinase K solution for 8 min, washed with PBS for 5 min, fixed with Z-fix for 5 min, washed with PBS for 5 min, incubated with rTDT enzyme and nucleotide mix in equilibration buffer for two hours, washed with 2× SSC for 5 min, counterstained with 2.5 m Hoechst 33342 in TBS for 10 min, and rinsed with TBS for 5 min. Coverslips were then mounted using VectaShield Vibrance and imaged using an A1R confocal on a Nikon Ti2 microscope.

### 5.3.13 Galectin-3 inhibitor experiments

At baseline, animals were assessed by electroretinogram, spectral domain coherence tomography (SD-OCT), and confocal scanning laser ophthalmoscope (CSLO) to evaluate any inherent structural or functional features or defects. Animals were injected with 15mg/kg of TD139 (33DFTG, catalog # AOB37408, AOBIOUS, Inc. Scranton, Pennsylvania) intraperitoneally daily beginning one day before light damage administration until day five post damage. Animals were then assessed using the same in vivo measures for retina architecture and structure changes.

### 5.3.14 Gene expression analysis (digital drop PCR)

Eyes were collected between 10 AM and 2 PM to standardize gene expression. The cornea and iris were removed via an incision, followed by the lens, and the neuroretina was separated from the RPE/choroid eye cup. Retinas were flash-frozen in RNase-free tubes and pre-chilled on dry ice. RPE/choroid eye cups were incubated in RNAprotect® Cell Reagent (Qiagen, Cat # 76106, Germantown, Maryland). for 10 minutes, with occasional agitation to release RPE cells. Cells were pelleted by centrifugation (>12,000 x g for 5 minutes), the supernatant was discarded, and the cells were stored at −80°C. RNA extraction was performed using the Qiagen RNeasy Mini Kit (Cat #74106). Samples were homogenized in RLT buffer with a stainless-steel bead, followed by ethanol addition and vortexing. The mixture was processed through an RNeasy column, washed with RW1 and RPE buffers, and eluted with nuclease-free water. The final RNA samples were stored at −80°C. cDNA synthesis was conducted using the Qiagen Quantitect RT kit.

#### Digital drop PCR (ddPCR) Reactions

Reaction mixes containing reverse transcriptase, primers, RT buffer, and QX200TM ddPCR EvaGreen Supermix (Bio-Rad: 186–4034) were added to 2μL of cDNA template for a total volume of 20 μL /well on the plate Twin-Tec plate (CAS # 951020320; Eppendorf, Enfield, CT). Fill empty well with RT Buffer and seal plate with tape film and spin down and mix. Plates were preheated at 95 C for 2 min/cycle. After using the droplet generator to generate droplets on the ddPCR plate, seal the droplet plate with foil film using the Biorad program. Then place the sealed Twin-Tec plate into ddPCR apparatus (QX200 Droplet Digital PCR (ddPCR™) System – Bio-Rad) and run the program as detailed in manufacturer’s manual.

### 5.3.15 Imaris analysis

The intensity, size, and distribution of Galectin-3 positive immune cells were analyzed using Imaris software 10.1.0 by Bitplane. Maximum intensity projection images of each RPE flat mount were processed using IMARIS 10.1.0 (Bitplane, Inc.), in which individual cells were identified, segmented, and quantified morphologically. Before converting and uploading images to Imaris, the corneal flaps and optic nerve heads were removed via the crop tool in Photoshop. Subretinal immune cell counts were conducted using the spots function in Imaris (artifacts and cell particulates were manually rejected) so that only cells with intact soma were quantified. Cell counts were normalized against double-blind manual cell counts of the same samples.

### 5.3.16 Statistical analysis

Statistical analysis was conducted using Prism 9.1.0 (on Mac OS X 14 Sonoma) (GraphPad Software, Inc., La Jolla, CA, USA). Data are presented as mean +/- standard deviation (SD), with statistical testing for individual datasets described in the Figure legends. A p-value <0.05 was considered statistically significant. Demographic distributions and sample sizes are summarized in Table 1. All statistical tests used are detailed in the <u>Figure Legends</u>.

## 5.4 RESULTS

### 5.4.1 Figure 1: Loss of PEDF is a Phenotype Modifier for Sensitivity to Phototoxic Damage in C57BL/6J

Expression of PEDF protects neurons and photoreceptors^26,45,46^. Conversely, loss of PEDF is linked to neurodegenerative disease phenotypes, including an autosomal dominant retinitis pigmentosa locus in human studies^24,47^. To determine if loss of PEDF sensitizes C57BL/6J mice to phototoxic damage, we crossed PEDF-null mice to mice with a hypomorphic mutation in the RPE65 gene, resulting in reduced sensitivity to light damage. We exposed these animals to 50,000 lux of light for 5 hours. We found that PEDF-null animals had more mottling in the fundus after LIRD than wildtype controls and experienced more retinal degeneration and thinning (see Figure. 1E-F). We quantified these changes amongst PEDF^+/+^, PEDF ^+/-^, and PEDF ^KO/KO^. We found that PEDF ^+/-^ behaved very similarly to PEDF ^+/+^ animals and showed minimal perturbances to ocular structure after LIRD (Fig. 1G-H). However, PEDF ^KO/KO^ showed significant losses of photoreceptor thickness and total retinal thickness compared to PEDF^+/+^ and PEDF^+/-^ animals (Figure 1G-H). Analysis: One-way ANOVA with Brown-Forsythe test and Barlett’s correction. Retinal thickness: PEDF^+/+^ vs. PEDF ^+/-^ p-value= not significant(ns); PEDF^+/+^ vs. PEDF ^KO/KO^ **p-value<0.01; PEDF ^+/-^ vs. PEDF ^KO/KO^ **p-value0.01. Photoreceptor thickness: PEDF^+/+^ vs. PEDF ^+/-^ = ns; PEDF ^+/+^ vs. PEDF ^KO/KO^ ****p-value<0.0001; PEDF ^+/-^ vs. PEDF ^KO/KO^ ****p-value<0.000. PEDF ^+/+^ n=5, PEDF ^+/-^n=4, PEDF ^KO/KO^ n=4). This data suggests that PEDF is protective against increased phototoxic damage.

**Figure 1:**
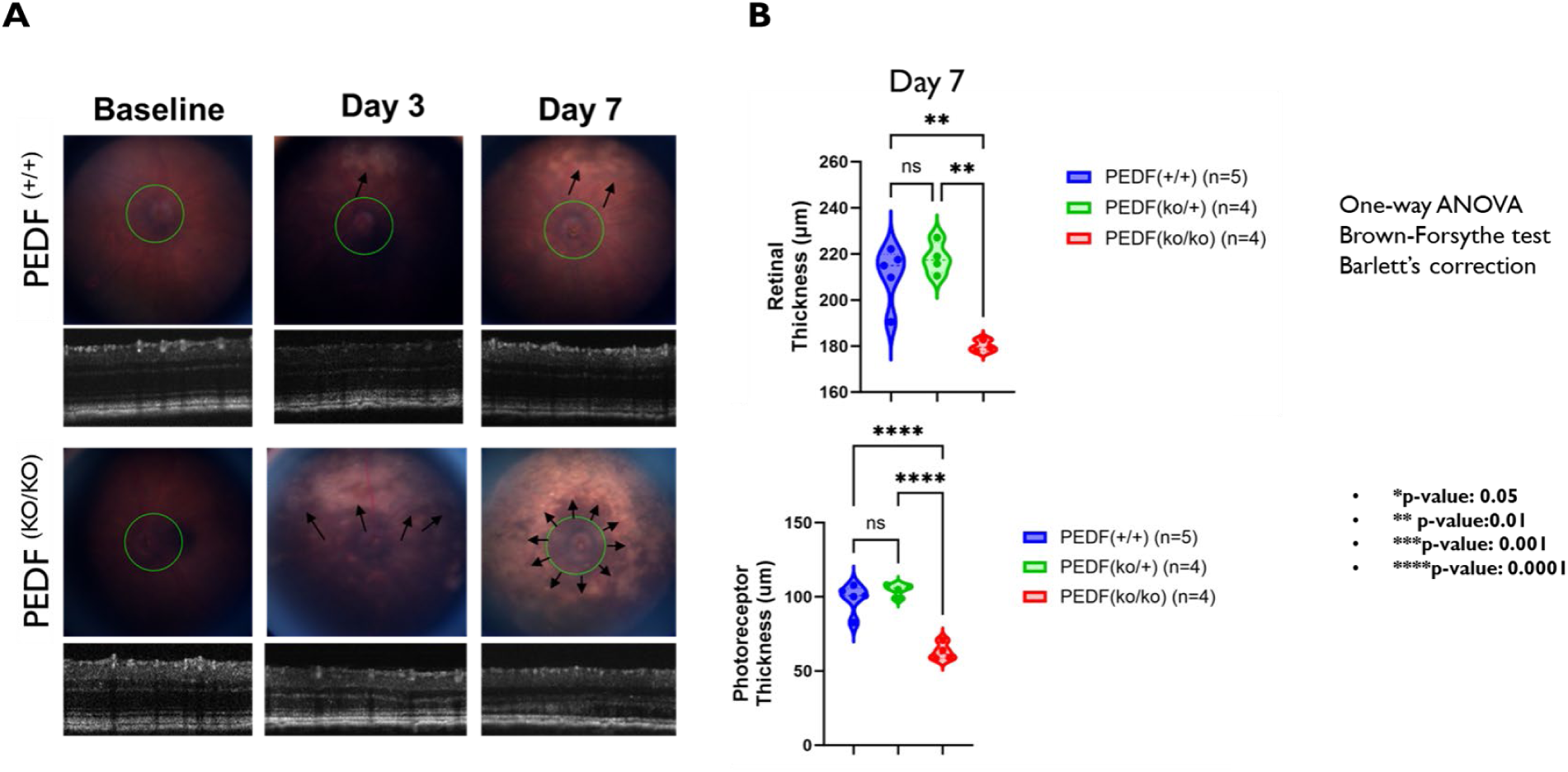
Loss of Pigment Epithelium Derived Factor Modifies Sensitivity to Phototoxic Damage in C57BL/6J Animals. Fundus and retinal C57BL/6J animals that express wild-type PEDF (PEDF +/+) or PEDF knockout (PEDF KO/KO) animals are shown. Figure 1A shows Spectral Domain Optical Coherence Tomography (SD-OCT) images of the Fundus and circular B-scans of the retinal architecture around the optic nerve. Top row: PEDF +/+ animals are shown in the top row at both baselines and on day seven post-LIRD. Bottom row: PEDF KO/KO animals at baseline and Day 7 Post LIRD. White arrows denote regions of damage-associated mottling of the fundus. Figure 1B shows the quantification of total retinal thickness and the thickness of the photoreceptor layer of PEDF +/+ n=5, PEDF KO/+ n=4, PEDF KO/KO= n=4 at Day 7 Post LIRD. One-way ANOVA Brown-Forsythe test with Barlett’s correction. * p-value< 0.05, ** p-value<0.01, *** p-value<0.001, **** p-value<0.0001

### 5.4.2 Figure 2: Loss of PEDF increases damage-associated autofluorescent dots at the level of the RPE

We used cSLO to capture dynamic changes at the level of the photoreceptor-RPE interface. At baseline, there were no differences or abnormalities between PEDF ^+/+^ (2A-B) or PEDF ^KO/KO^ (2F- G) in the vasculature or at the level of the RPE interface. However, when assessing the same animals on Day 7, the number of damage-associated punctate at the RPE-photoreceptor layer was significantly increased in the PEDF ^KO/KO^(2H-J) animals compared to the PEDF ^+/+^ (2C-E). This data suggests that PEDF-null animals have improved response to damage via the appearance of damage-associated foci at the RPE-photoreceptor interface.

**Figure 2:**
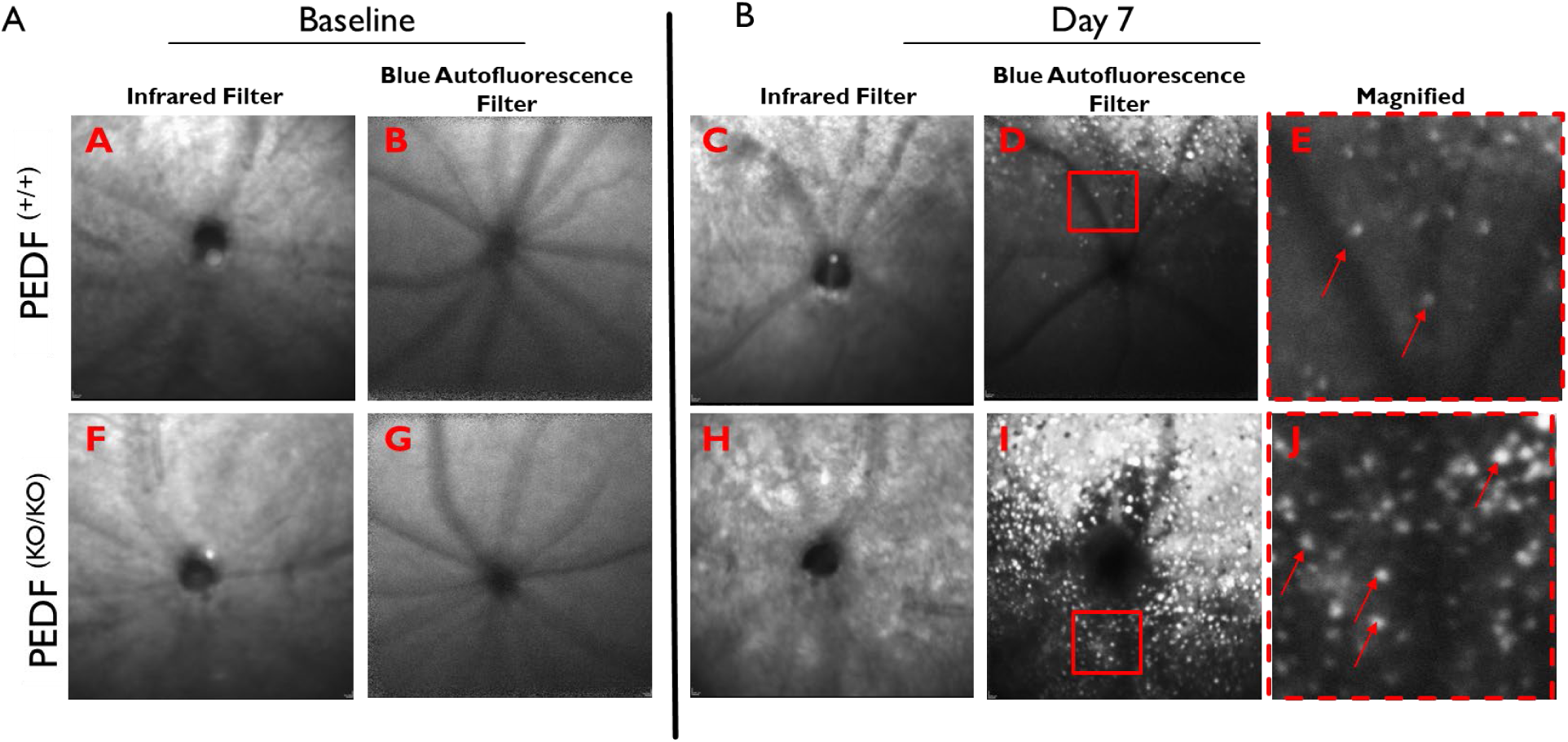
Loss of PEDF Increases Damaged-Associated Autofluorescent Dots at the Level of the RPE. Heidelberg Spectralis cSLO (confocal scanning laser ophthalmoscope) images show an increased accumulation of autofluorescent dots/granules at the level of the photoreceptor-RPE interface after phototoxic damage that is not present at baseline. Images were taken at −12 diopters at the level of the interdigitations of RPE and photoreceptors using both the infrared (to detect vascular architecture) and the blue autofluorescence filter (to detect fluorescent dots). Representative Images at baseline for PEDF ^+/+^(2A-B) and PEDF ^KO/KO^ (2F-G) and at Day 7 Post LIRD (PEDF ^+/+^: 2C-E; PEDF ^KO/KO^: 2H-J). A Zoom (red box) of each representative image with red arrows highlighting individual dots.

### 5.4.3 Figure 3A: There is regionality to the damage phenotype in PEDF knockouts compared to the wild type.

We used H&E to quantify the number of nuclei remaining in the outer nuclear layer (ONL) after LIRD damage to assess the degree of the damage and morphological changes. PEDF ^+/+^ animals still had relatively normal morphology with intact RPE layer and photoreceptor inner and outer segments before and after LIRD (Figure 3A-B). However, the PEDF ^KO/KO^ animal displayed a significant loss of total retinal thickness, a drastically diminished ONL, an almost complete loss of photoreceptor inner and outer segments, and compromised RPE integrity (shown via white arrows: differences in RPE thickness; Fig. 3C-D). There were regional characteristics to this damage phenotype in the PEDF ^KO/KO^ animals, with retinal structures on the superior portion of the eye being more severely diminished compared to the inferior region of the eye (Fig.3E). A similar phenotype was also shown in day five after damage [data not shown]. (Analysis: One-way ANOVA with Brown-Forsythe test and Barlett’s correction; ^##^ p-value<0.01 and ^###^p-value< 0.001; PEDF ^+/+^ n=4, PEDF ^KO/KO^ n=4). This phenomenon is characteristic of light damage models, as described by Rapp and Williams^48,49^ and our data confirms that.

**Figure 3:**
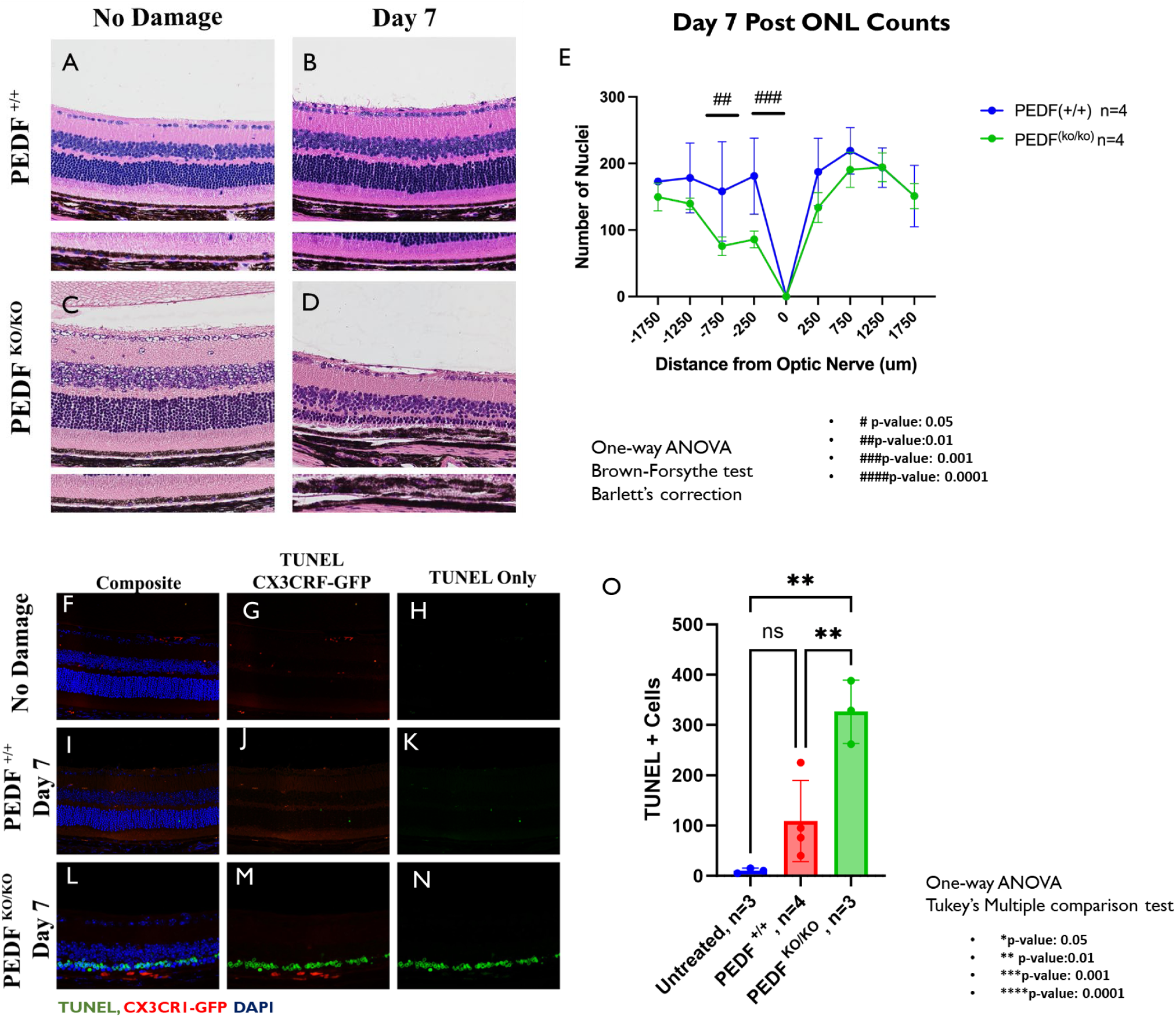
Loss of PEDF Results in Regional Damage and Increases Apoptosis of Photoreceptor Cells. The morphology of the postmortem tissue shows significant regional alterations in retinal architecture. Figure 3A-B shows a representative image of PEDF ^+/+^ with no damage and day seven post-LIRD. Representative images of PEDF ^KO/KO^ animals with no damage(Figure 2C) and day seven post-damage (Figure 2D) are shown. Figure 2D shows severe loss of the outer nuclear layer (ONL), disruption of the photoreceptor inner and outer segment layer, and aberrations in the RPE monolayer in PEDF ^KO/KO^ compared to PEDF ^+/+^ controls at day five post-light damage. Figure 3E quantifies ONL counts from - 1750 microns(superior) to 1750 microns (inferior) on either side of the optic nerve. The damage is regionally isolated to the superior portion of the retina and is significantly between PEDF KO/KO n=4 and PEDF ^+/+^ n=4. One-way ANOVA with Brown-Forsythe test and Barlett’s correction. # p-value<0.05, ## p-value<0.01, ### p-value<0.001, #### p-value <0.0001. The loss trend was the same on day seven post-LIRD (data not shown). Figure 3F-N shows representative images of retinal sections stained for TUNEL (green), immune cells via CX3CR1-GFP (red), and cell nuclei (DAPI) of no damage control (3F-H), Day 7 PEDF ^+/+^(3I-K) and, Day 7 PEDF ^KO/KO^ (3L-N). These data are quantified in Figure 3O and show that PEDF ^KO/KO^ have significantly more TUNEL-positive cells than either the untreated (** p-value<0.01) or the PEDF ^+/+^ (** p-value<0.01) group.

Previous light studies in rats have suggested that peak DNA damage occurs within the first 8-16 hours after damage ^50^. To assess if PEDF ^KO/KO^ animals were still undergoing significant levels of active apoptosis at day 7, we stained for DNA fragmentation using TUNEL and immune cells using CX3CR1-GFP. PEDF ^KO/KO^ animals had significantly more apoptotic cells at day 7, resulting in a more depleted outer nuclear layer than wild-type controls. Additionally, there are more immune cells in the PEDF ^KO/KO^ subretinal space compared to the wild-type animals at the same time point (Fig. 3L-N; quantified in Fig. 3O: Analysis: One-way ANOVA with Tukey’s multiple comparison tests: untreated vs. PEDF^+/+^ p-value=ns; untreated vs. PEDF ^KO/KO^ **p-value <0.01; PEDF ^+/+^ vs PEDF ^KO/KO^ **p-value<0.01. untreated n=3, PEDF ^+/+^ n=4, PEDF ^KO/KO^ n=3.) This data suggests that loss of PEDF increased regional loss of photoreceptors after light damage.

### 5.4.4 Figure 4: PEDF KO/KO animals’ RPE fails to increase rhodopsin metabolism after light damage.

Loss of PEDF in the RPE affects aging and RPE functional deficiency^2,51^. To assess changes in RPE function in the absence of PEDF, we performed a rhodopsin metabolism assay as a proxy for RPE phagocytic capacity, a critical function of the RPE. We found that at day seven after LIRD, PEDF ^+/+^ animals significantly increased rhodopsin metabolism in response to damage. However, PEDF ^KO/KO^ mice failed to significantly increase rhodopsin metabolism, although they showed increased damage compared to wild-type littermate controls (See Figure 4F; quantified in Fig. 4G: Two-way ANOVA with Tukey’s multiple comparison test, *p-value<0.05). Defects in phagocytosis of PEDF ^KO/KO^ mice have been previously documented^10^. These data suggest that loss of PEDF results in reduced capacity for phagocytosis by the RPE.

**Figure 4:**
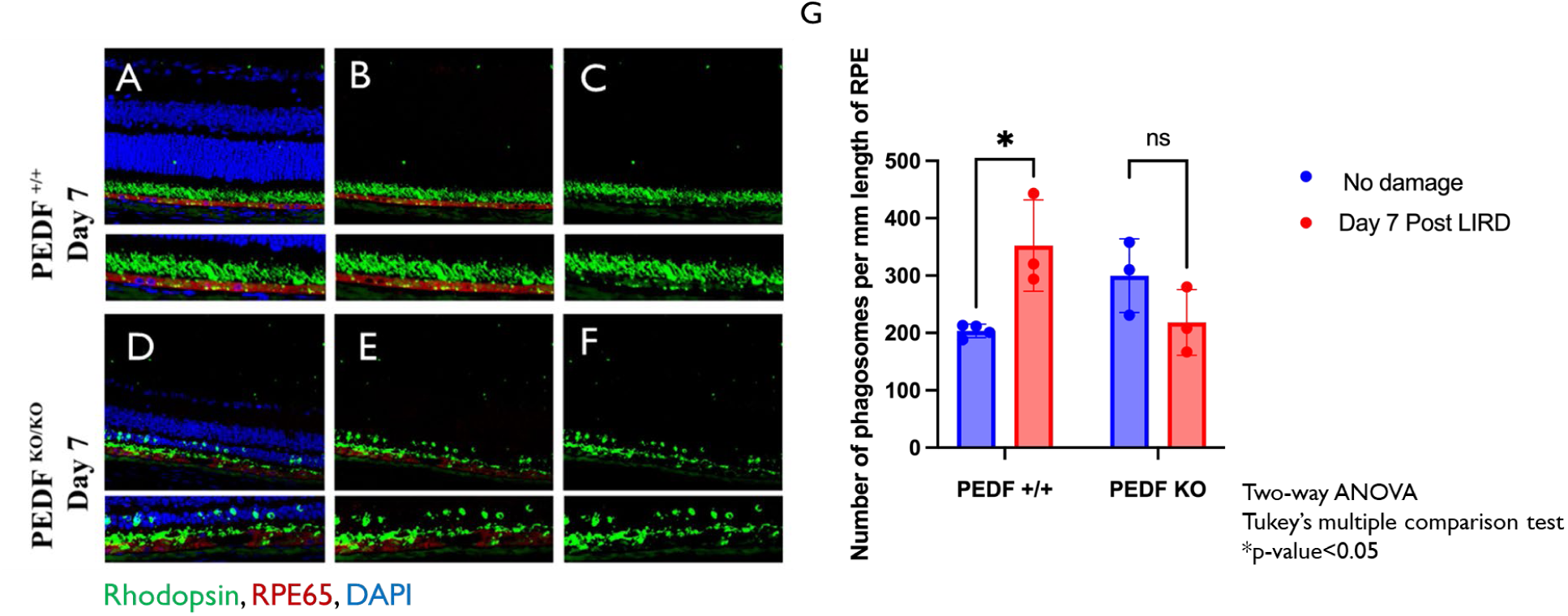
PEDF ^KO/KO^ RPE Fail to Increase Rhodopsin Metabolism after Light Damage. Loss of PEDF results in a suboptimal production of phagosomes by the RPE after light-induced retinal damage. Figure 4A-F shows representative retinal immunofluorescence images of a PEDF ^+/+^ and PEDF ^KO/KO^ at day 7 Post-light damage. The sections were stained with Rhodopsin(green) to visualize shed rod outer segments and phagosomes, Best1(red) was used to visualize the RPE monolayer, and cell nuclei were stained with DAPI (blue). Figure 4G, notably, the PEDF ^+/+^ animals significantly increase production to redress clearance demands at day seven post-LIRD compared to untreated PEDF^+/+^(Two-way ANOVA, Tukey’s multiple comparison test, *p-value<0.05). However, while PEDF ^KO/KO^ animals had a more significant accumulation of phagosomes at baseline, they failed to increase phagosome production after light damage.

### 5.4.5 Figure 5: PEDFKO/KO results in loss of retinal function following light stress

We also assessed for functional changes using electroretinograms to accompany the distinctive in vivo and post-mortem histology analysis that we performed. Under scotopic conditions, we found that at baseline until three days post-LIRD, there was no significant difference between genotypes in either a- or b-wave function. However, by days 5 and 7, there were significant defects in a- and b-wave amplitudes of PEDF ^KO/KO^ compared to wild-type littermates (Fig. 3A-B: Two-way ANOVA with Sidak’s Multiple comparison correction. Scotopic a-wave-Day 5: PEDF ^+/+^ vs. PEDF ^KO/KO^ **-p-value<0.01. Day 7: **p-value<0.01 n=3-7/group/timepoint. Scotopic b-wave: Day 5: *p-value<0.05. Day 7: *p-value<0.05). To accompany the rhodopsin metabolism analysis, we used c-wave analysis as a proxy to evaluate the RPE function. We found that after light damage, there is not a significant difference between PEDF ^+/+^ and PEDF ^KO/KO^ animals until day seven post-LIRD damage (Fig5.C: Two-way ANOVA with Sidak’s multiple comparison correction: PEDF^+/+^ vs. PEDF ^KO/KO^; Day 5-ns; Day 7 *p-value<0.05). This datum aligns with the functional deficits observed in the RPE in our immunofluorescence data from Figure 4. We also assessed the scotopic and photopic waveforms of PEDF ^KO/KO^ compared to PEDF ^+/+^ at baseline and day seven post-LIRD. PEDF ^KO/KO^ animals have a slightly lower b-wave and c-wave amplitude compared to PEDF ^+/+^ littermate controls at baseline (Fig.5D); however, there were no defects in phototopic function (Fig. 5F). At day seven after damage, both scotopic and photopic waveforms worsened in PEDF ^KO/KO^ animals compared to PEDF ^+/+^ animals (Fig. 5E and 5G). These data suggest that the loss of PEDF negatively affects the retina and RPE function and leads to increased damage after LIRD compared to PEDF ^+/+^ littermates.

**Figure 5.**
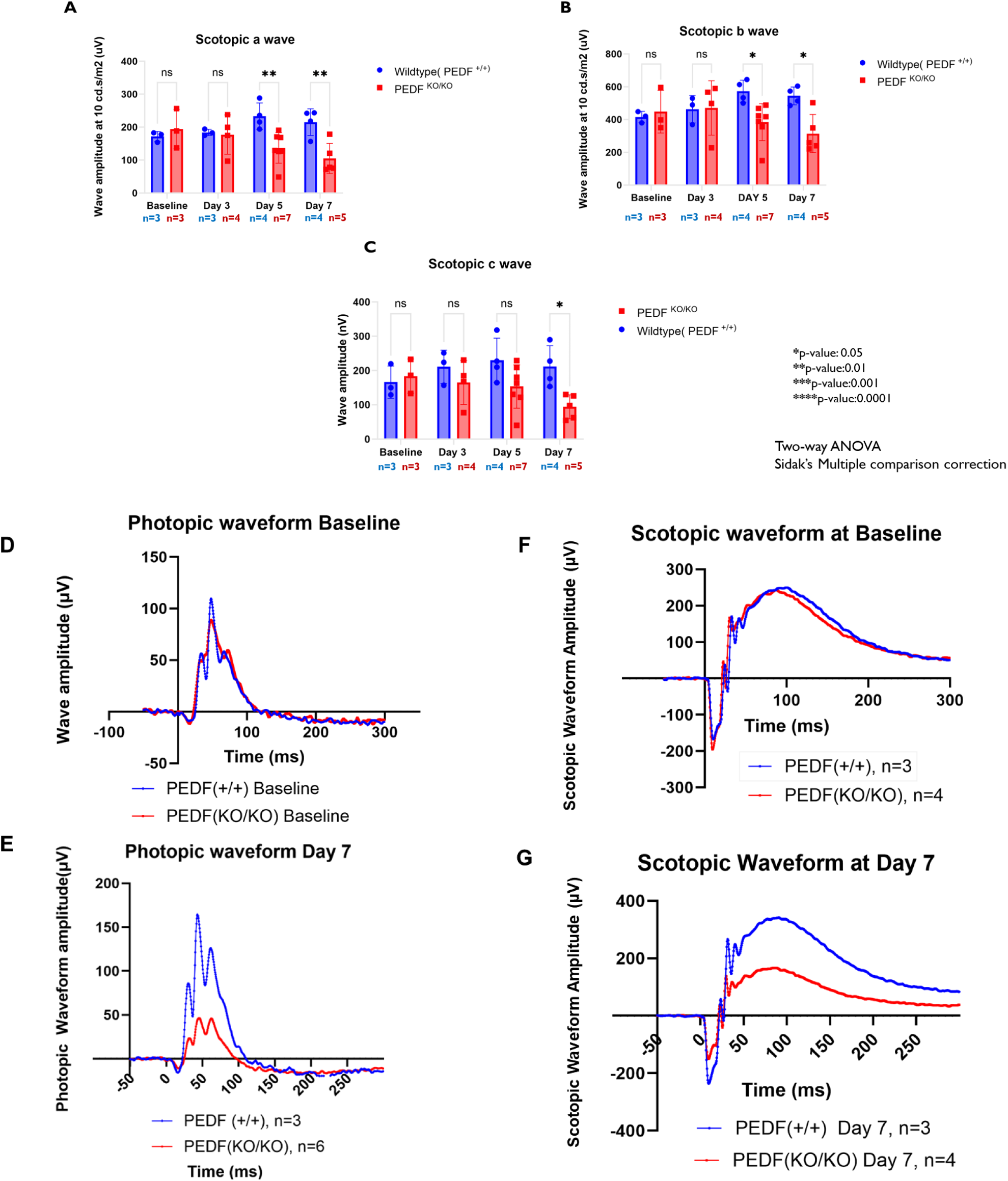
The Loss of PEDF leads to significant deficits in visual function after light damage exposure. The figure shows the maximal visual output of *a*-wave, b-wave, and *c*-wave at a flash intensity of 10 candelas/second/meters^2^ (cd.s/m^2^). These data show no statistically significant difference in the visual function of the PEDF ^KO/KO^ compared to PEDF ^+/+^ at baseline or on day three after light damage. However, after day three there is a notable decrease in visual function of PEDF ^KO/KO^ animals in both a- and b-wave amplitudes at 10Hz that persists to day 7(a-wave: day 5: ** p-value<0.01; day 7: ** p-value<0.01 and b-wave: day 5: *p-value <0.05; day 7: *p-value<0.05. n=3-7/time point/group) see Figure 5A and 5B; Two-way ANOVA with Sidak’s multiple comparison correction). Significant loss of the c-wave amplitudes is delayed to day seven post-light damage (See Fig. 2C: Two-way ANOVA with Sidak’s multiple comparison corrections, day 7: * p-value<0.01). The scotopic waveforms of PEDF KO/KO mice also reveal a slight depression in the waveform amplitude at baseline compared to PEDF ^+/+^ (n=3-4/genotype). This reduction in waveform amplitude is more pronounced at day seven post-LIRD (n=5/genotype; See Figures 5D and 5E). Photopic waveforms show a similar trend as scotopic waveforms with significantly reduced amplitudes in PEDF KO/KO at day 7 compared to PEDF ^+/+^ littermates (See Fig. 5F-G).

### 5.4.6 Figure 6: PEDF ^KO/KO^ Results in Suppression of the Damaged-Associated Increase in IGF1 Expression after Light Damage

Studies of hypoxic trauma, diabetic retinopathy, and pharmacological damage in the eye have linked the expression of PEDF and insulin-like growth factor 1(IGF-1) to the protection of RPE cells and other ocular structures after insult ^52–54^. To determine if loss of PEDF impacts the expression of IGF-1 after damage, we used immunofluorescence to stain retinal sections of PEDF ^+/+^ and PEDF ^KO/KO^ animals. We quantified the expression of IGF-1 from baseline until day seven post-damage. Notably, PEDF ^KO/KO^ animals showed significant reductions in IGF-1 starting at day three compared to wildtype littermates ( Fig 6Q: Two-way ANOVA with Tukey’s multiple comparison test, n=3-4 animals/group/timepoint. Day 3: ****p-value<0.0001; Day 5: ****<0.0001; Day 7: ****p-value<0.0001). Increased infiltrating galectin-3 positive immune cells were found at the RPE-photoreceptor interface in PEDF ^KO/KO^ animals and significantly more damage via loss of ONL thickness compared to wildtype littermates (See Fig. 6A-P). To confirm these findings, we tested the protein expression of IGF-1 in PEDF ^+/+^ and PEDF ^KO/KO^ animals. At baseline, there is no significant difference in IGF-1 expression among PEDF ^+/+^ and PEDF ^KO/KO^ animals (Two-way ANOVA with Tukey’s multiple comparison test. N=3-6 animals/group/timepoint. Baseline: PEDF +/+ vs. PEDF KO/KO =ns). PEDF ^+/+^ animals significantly increased IGF-1 expression by day seven after damage (PEDF +/+ no damage vs. PEDF ^+/+^ Day 7 post *p-value<0.05). Notably, the expression of IGF-1 in response to damage was significantly dampened in PEDF ^KO/KO^ compared to PEDF ^+/+^ animals at day 7 (PEDF ^+/+^ Day 7 vs. PEDF ^KO/KO^ Day 7 **p-value<0.01). Immune cells, like microglia, with high expression of IGF1 are associated with neuroprotection ^55,56^. We found that subretinal immune cells in the PEDF ^+/+^ animals on day 7 showed a prominent expression of IGF1 in the cell body/cytoplasm. However, the subretinal immune cells in the PEDF ^KO/KO^ had very little to no expression of IGF-1. These data may suggest that loss of PEDF results in global loss of IGF-1 expression and increased recruitment of IGF-1 deficient immune cells.

**Figure 6:**
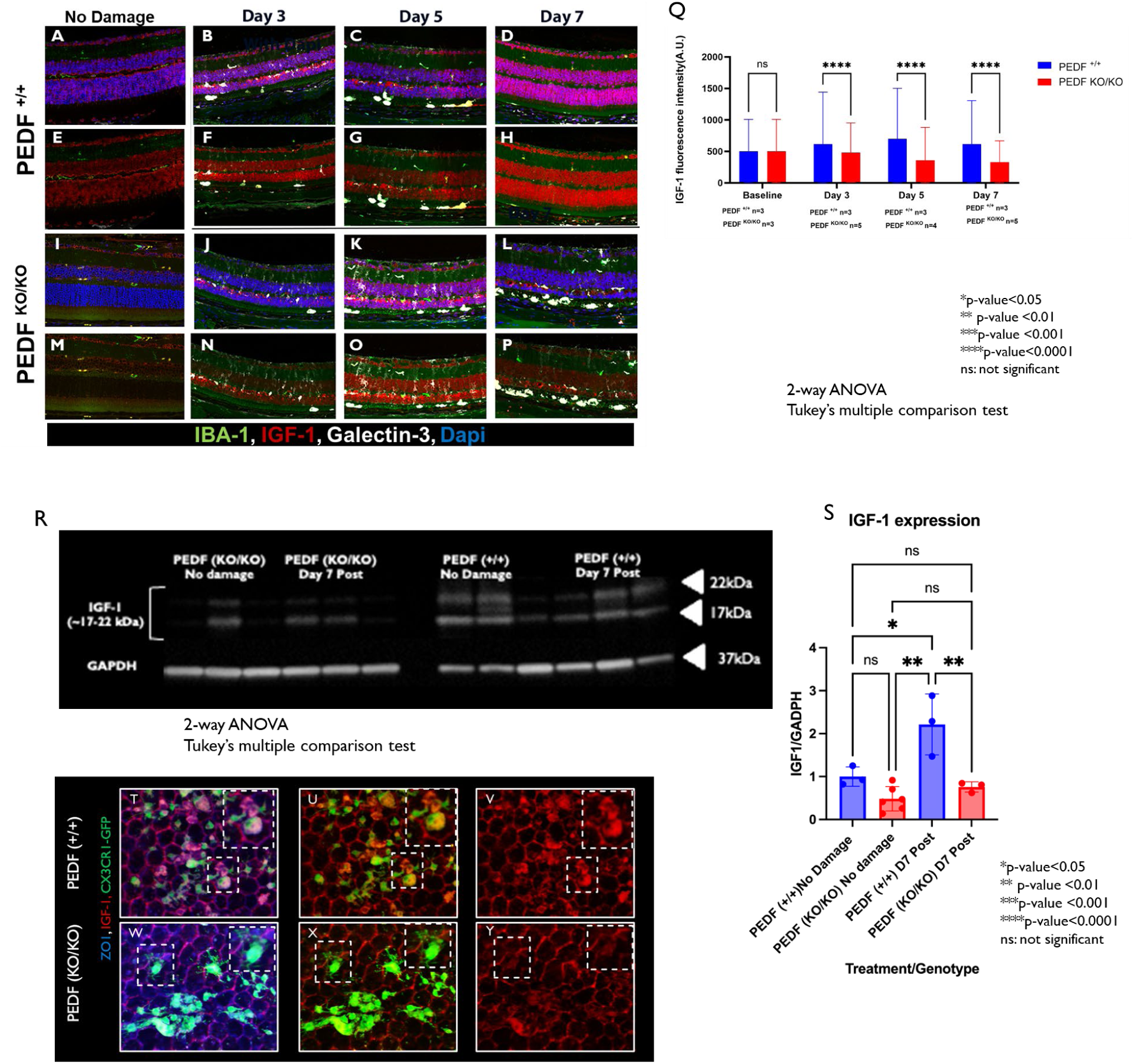
Loss of PEDF Suppresses the Damage-Associated Increase in IGF-1 after Light Damage. Retinal sections were collected at days 3, 5, and 7 post light damage and stained for the neurotrophic factor, IGF-1(red), the immune cell marker, IBA1(green), immune cell activation marker, Galectin-3(white), and dapi(blue). Staining showed that at baseline, there was no significant difference between PEDF ^+/+^ and PEDF ^KO/KO^ animals without damage. Figure 6A-P is a representative image showing the degree of expression of IGF-1 and Galectin-3 and the infiltration of immune cells from the retina to the subretinal space. After light damage, there is an increase in Galectin-3 positive cell expression in both genotypes at day 3, with the earliest deposition at the photoreceptor-RPE interface occurring at day 3. By day 7, only the PEDF ^KO/KO^ animals still have Galectin-3 positive cells at the interface of the photoreceptors-RPE. Additionally, when quantifying the immunofluorescent signal of IGF-1, there are statically significant differences between the PEDF ^+/+^ and PEDF ^KO/KO^ as early as day 3. The levels of IGF-1 continue to decrease until day seven post LIRD (see Fig. 6Q). Analysis: Two-way ANOVA with Tukey’s multiple comparison test, n=3-5/ group/time point. * p-value< 0.05, ** p-value<0.01, *** p-value<0.001, **** p-value<0.0001. In Figure 6R, we confirm this finding via total eye cup expression of IGF-1 normalized to GAPDH in no damage controls versus at day seven post-LIRD via western blot. Figure 6R quantifies the total expression of IGF-1 between PEDF ^+/+^ and PEDF ^KO/KO^ before and after LIRD. Analysis: Two-way ANOVA with Tukey’s multiple comparison test, n=3-6/group/timepoint. Subretinal immune cells recruited to RPE in PEDF ^KO/KO^ have lower expression of IGF-1 than PEDF +/+ animals. Figure 6T-Y shows a representative image of PEDF ^+/+^( 6T-V) and PEDF ^KO/KO^ (6W-Y) stained for ZO1(blue), IGF-1(red), and CX3CR1-GFP (green) to look for heterogeneity in the immune cell population.

### 5.4.7 Figure 7: Loss of PEDF results in robust inflammatory response compared to wildtype controls

Pigment epithelium-derived factor regulates inflammatory responses in multiple diseases, including diabetic retinopathy, dry eye disease, and cancer studies ^17,21,57–61^. Specifically, the 44-mer and 17-mer PEDF peptides have been associated with antagonizing IL-6 production, thus suppressing chorioretinal inflammation ^62^. We used immunofluorescence staining of RPE flat mounts to evaluate how the loss of PEDF affects the recruitment of subretinal immune cells at different time points after LIRD. The number of subretinal immune cells in PEDF ^KO/KO^ and wildtype littermates is comparable at baseline. However, after LIRD, PEDF ^KO/KO^ animals had significantly more recruitment of subretinal immune cells by day five than wildtype littermates (See Fig. 7A-D; quantified in Fig. 7E: Two-way ANOVA with Sidak’s multiple comparison test, Day 5: PEDF ^+/+^ vs. PEDF ^KO/KO^ **p-value 0.01). The number of subretinal immune cells peaked on day 7 (****p-value 0.0001). Additionally, the cells had higher expression of galectin-3, a pleiotropic, β-galactoside-binding protein associated with reactive microglia, compared to wildtype littermate controls at day seven post ^33^.

**Figure 7:**
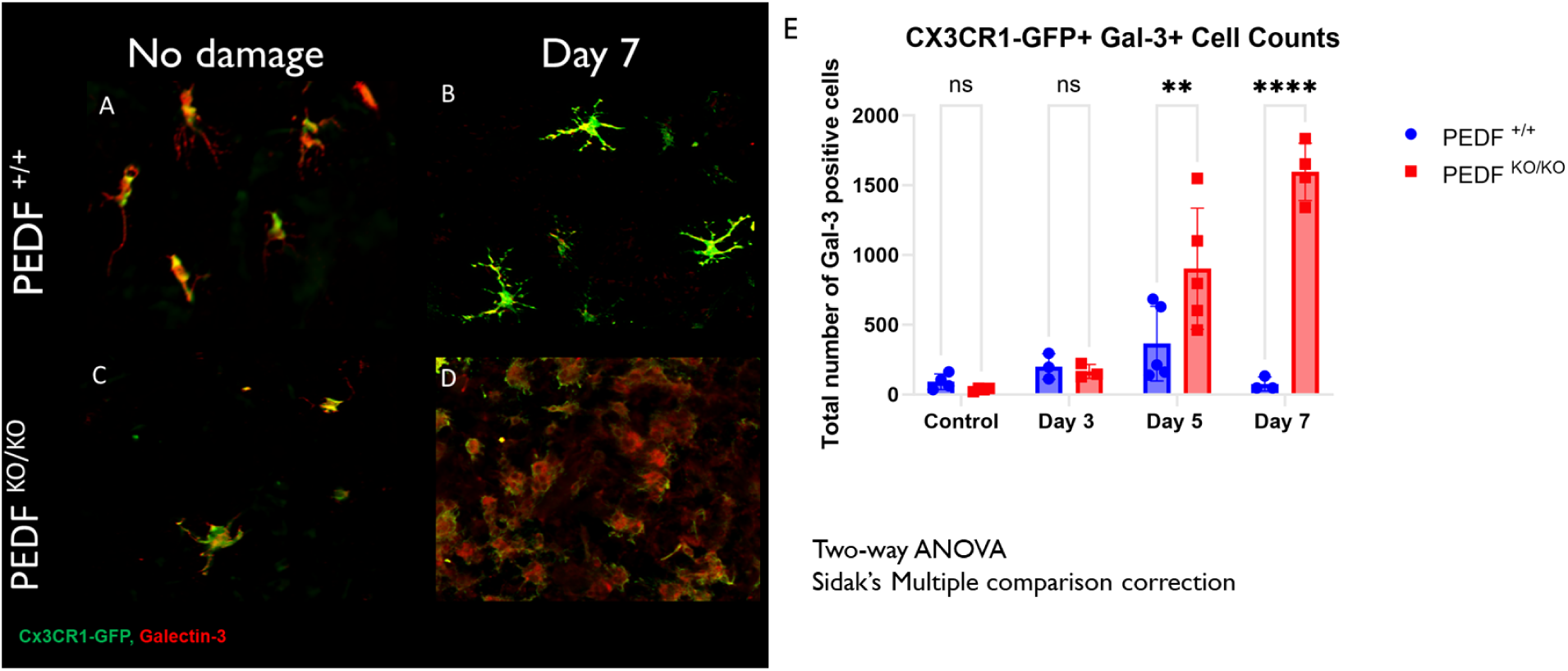
Loss of PEDF increases infiltration of galectin-3^+^ immune cells compared to PEDF ^+/+^. We collected RPE flat mounts to assess if PEDF ^KO/KO^ animals showed an increased inflammatory profile and stained them for Galectin-3 (red) and CX3CR1-GFP(Green). We found that PEDF ^KO/KO^ animals were like PEDF ^+/+^ animals at baseline and up to day three post-LIRD damage. However, by day 5, there was the inflammation phenotype significantly increased in PEDF ^KO/KO^ animals compared to littermate controls * p-value< 0.05, ** p-value<0.01, *** p-value<0.001, **** p-value<0.0001 (Analysis: Two-way ANOVA with sidak’s multiple comparison correction. N=3-5 animals group/ time point. p-value: Day 5: ** vs Day 7 **). Figure 7A-D shows a representative image of the subretinal immune cell morphology in PEDF ^+/+^ and PEDF ^KO/KO^ animals at baseline and Day 7. Figure 7E shows the total number of Gal-3 positive cells counted from baseline to day seven post-LIRD between PEDF ^+/+^ and PEDF ^KO/KO^

### 5.4.8 Figure 8: Loss of PEDF differentially affects Lgals and Nlrp3 gene expression

To determine if loss of PEDF differential affects inflammasome activation after LIRD, we first used digital drop PCR to assess mRNA expression of both *Lgals3* and *Nlrp3* in both the retina (data not shown) and RPE. Lgals3, the gene that encodes galectin-3, mRNA expression was significantly lower in the RPE of PEDF ^KO/KO^ animals compared to wildtype littermate controls at baseline (Two-way ANOVA with Tukey’s multiple comparison test. *p-value<0.05). However, the amount of the transcript significantly increases on day 7 in PEDF ^KO/KO^ animals compared to wildtype littermates at the same time point (**p-value< 0.01). Additionally, Nlrp3 mRNA in the RPE only increased significantly at day seven post-LIRD in PEDF ^KO/KO^ compared to wildtype littermates (*p-value<0.05). The supplemental information can find the mRNA expression of LGALS3 and NLRP3 in RPE and SNAI1, IL-6, and IL1-beta expression in retina and RPE. The loss of PEDF differentially regulates genes that encode galectin-3 and inflammasome-associated protein, Nlrp3, at baseline and after LIRD, implicating PEDF in regulating galectin-3 gene expression.

### 5.4.9 Figure 9: Loss of PEDF reduces total Galectin-3 expression

Previous studies have identified immune cells recruited to the subretinal space as a unique subset enriched for galectin-3 ^63,64^. To investigate the relationship between the loss of PEDF and galectin-3 expression, we performed protein expression analysis via western blot at baseline and day seven post-LIRD in PEDF ^KO/KO^ compared to PEDF ^+/+^. PEDF ^KO/KO^ animals, at baseline, had significantly lower galectin-3 protein expression than those of PEDF ^+/+^ littermate controls (PEDF ^+/+^ vs. PEDF ^KO/KO^ Baseline ****p-value<0.0001). This data substantiated results from Figure 8A, which showed lower Lgals3 mRNA expression in PEDF ^KO/KO^ animals at baseline. However, while the level of galectin-3 protein expression in PEDF ^KO/KO^ animals increases after phototoxic damage, it remains suboptimal to PEDF ^+/+^ animals at the same time point (Two-way ANOVA with Tukey multiple comparison test, n=3/group/timepoint. PEDF ^+/+^ vs PEDF ^KO/KO^ Day 7 ***p-value 0.001). These data suggest the loss of PEDF significantly affects the protein expression of Galectin-3 both before and after LIRD.

**Figure 8:**
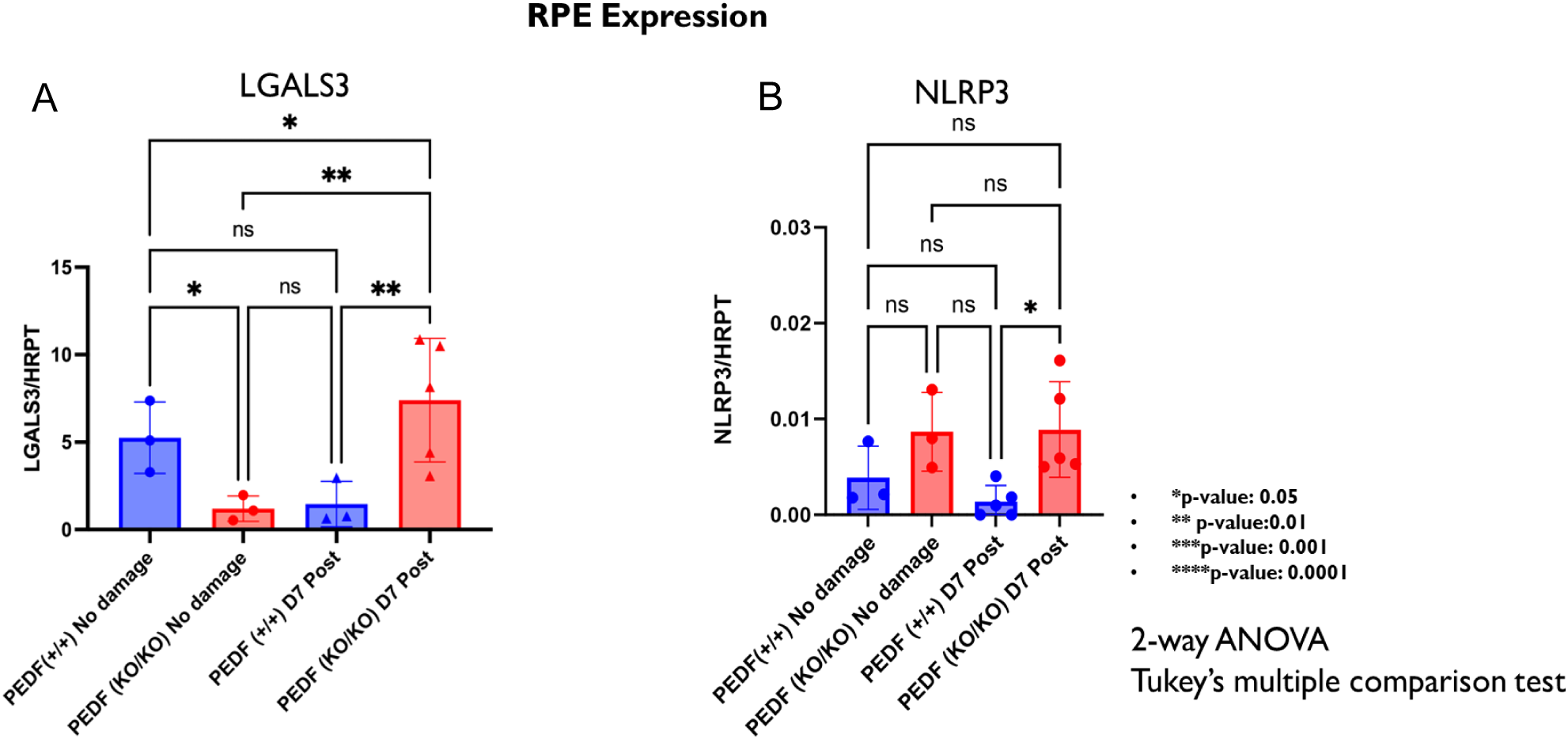
Loss of PEDF Increases Galectin-3 Gene Expression at Day 7 Post LIRD Compared to Wildtype Littermates. Retinal and RPE tissues were collected separately, and RNA was extracted from each tissue sample type. Figure 9A quantifies Lgals3 and Nlrp3 gene expression normalized to HRPT in the retina between PEDF +/+ and PEDF ^KO/KO^ at baseline and Day 7 Post LIRD. Figure 9A-B shows the gene expression of Lgals3 and Nlrp3 at the same time points in the RPE. The Lgals3 expression in the RPE Two-way ANOVA; PEDF ^KO/KO^ baseline vs. PEDF KO/KO Day 7: *p-value<0.05; PEDF ^+/+^ Day 7 vs. PEDF ^KO/KO^ Day 7: *p-value<0.05. However, only at day 7 in the RPE is Nlrp3 expression significantly different in the PEDF ^KO/KO^ compared to littermate controls(*p-value<0.05)

**Figure 9:**
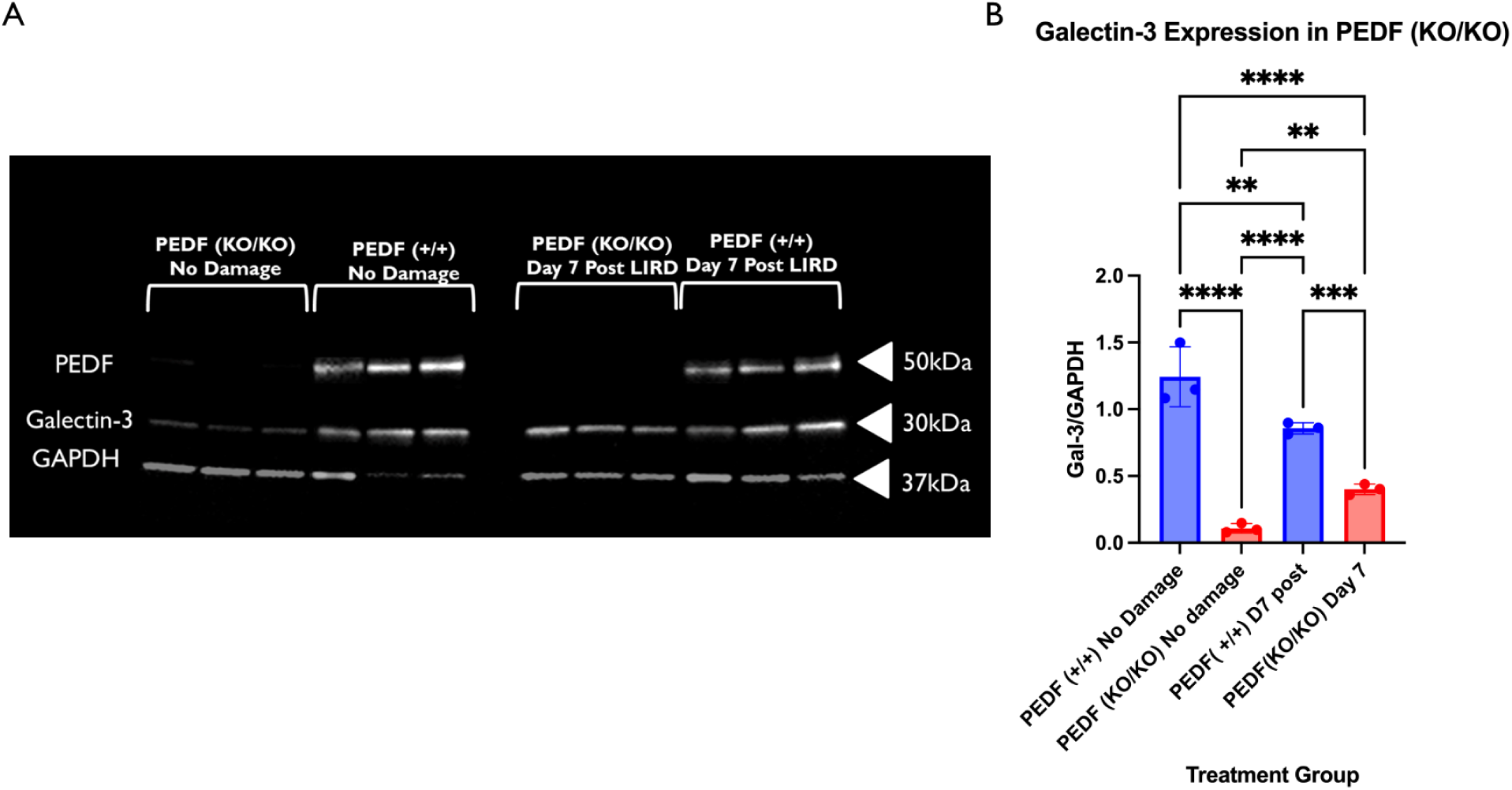
Loss of PEDF Reduces Total Galectin-3 Expression Before and After LIRD. PEDF ^KO/KO^ animals have significantly lower expression of Galectin-3 at baseline compared to littermate controls. Additionally, after damage, there is a suboptimal increase in Galectin-3 protein expression on day seven post-LIRD. PEDF ^+/+^ animals dampen galectin-3 expression in response to LIRD damage at day 7, suggesting differential temporal regulation of the protein when PEDF is present compared to when it is not. Figure 9A shows a western blot that was probed for PEDF (50kDa), Galectin-3(∼30kDa), and GAPDH (∼37kDa) loading control. The results from Figure 9A are quantified in Figure 9B and show that there are significant differences in Galectin-3 expression at both baselines (Two-way ANOVA with Tukey’s multiple comparison correction. **** p-value<0.0001. sample sizes: 3 animals/group/time point) and at Day 7 (****p-value<0.0001) between PEDF ^KO/KO^ and PEDF ^+/+^ animals. While Galectin-3 expression increases in the PEDF ^KO/KO^ animals at day seven compared to baseline, it is still dampened compared to the Gal-3 expression of PEDF ^+/+^ at the same time point.

### 5.4.10 Figure 10: Inhibition of Galectin-3 with TD139 significantly decreases PEDF levels after light damage

Previous studies have correlated increased expression of galectin-3 with poor clinical outcomes in multiple eye diseases ^65–70^. Additionally, the inhibition galectin-3 by genetic manipulation or pharmacological targeting dampened immune cell activity ^71^. To determine if dampening the galectin-3 expression would be protective after LIRD damage, we pharmacologically inhibited Galectin-3 in PEDF ^+/+^ animals using TD139 to determine if inhibiting galectin-3 was protective after LIRD. We found that treatment with galectin-3 inhibitor (TD139) did not significantly affect galectin-3 protein levels.

However, we did notice significant differences in the visual function of animals without LIRD exposure (data not shown). Interestingly, we found that animals treated with galectin-3 inhibitor had a worse damage phenotype than LIRD-only controls. Surprisingly, PEDF levels in animals treated with TD139 and LIRD were significantly lower than in the LIRD-only control group (One-way ANOVA with Tukey’s multiple comparison test. n=3 animals/group. PEDF ^+/+^ No damage vs. PEDF ^+/+^ LIRD only: p-value=ns; PEDF ^+/+^ no damage vs. PEDF ^+/+^ LIRD + Gal-3 inhibitor ***p-value<0.001; PEDF ^+/+^ LIRD only vs. PEDF ^+/+^ LIRD + Gal-3 inhibitor *p-value<0.01). Treatment with TD139 alone does not affect visual function or Galectin-3 protein expression compared to vehicle only(See Supplemental Figure 1). These data suggest a potential correlation between PEDF and Galectin-3 expression since inhibition of galectin-3 significantly decreases PEDF expression.

## 5.5 DISCUSSION

The findings from this study reveal that PEDF plays a significant regulatory role in facilitating immune privilege and suppressing inflammation to protect vulnerable tissues from damage within the ocular microenvironment. Previous studies have evaluated and purported the protective role of PEDF against photoreceptor death in albino rat models under various light damage conditions; these studies showed that intravitreal supplementation with exogenous PEDF was protective; however, the mechanism for this protection was not established^72,73^. These studies were limited in that they used albino animals, which are not as translatable to normal vision in humans, and they used This study aimed to examine the influence of PEDF on the outcome of visual function, galectin-3 positive subretinal immune cell recruitment, and effects on the neurotrophic factor, IGF-1, after light damage. By employing a global deletion model of PEDF and comparing the multiple visual metrics to wildtype controls, we could identify phenotypic shifts during damage resolution that coincide with expression changes in IGF-1 and Galectin-3. Studying these molecular mechanisms may be the basis for better understanding and predicting the pathological onset of disease, reveal new pathway interactions for conserved biomarkers, and present new considerations for therapeutic approaches employing gene therapy. To our knowledge, our study is the first to evaluate the potential regulatory axis of PEDF-Galectin-3-IGF-1 in visual function. Additionally, according to our understanding, this is the first study to implicate PEDF in the modulation of galectin-3 expression in the eye. Overall, our results implicate the loss of PEDF as an essential regulator of both IGF-1 and Galectin-3 expression after light damage, suggesting an additional level of RPE-mediated regulation of immunosuppression in the ocular microenvironment.

Immune privilege in the eye requires an intact RPE monolayer, which secretes factors that suppress the immune response, controls the maturation of immune cells, and leads to apoptosis of infiltrating macrophages, magnifying the role of RPE in facilitating immunomodulation^74–81^. Studies of pigment epithelia derived from various ocular tissues suggest that immunosuppression is achieved by cell-cell contact, soluble factors, or both 240, depending on the source of epithelia. The retinal pigment epithelia predominantly utilize secreted, soluble factors to suppress immune cell activation. Previous studies have described the immunomodulatory functions of the RPE via the secretion of cytokines and neuropeptides, like alpha-macrophage stimulating hormone(⍺-MSH) and Neuropeptide Y(NPY)^79,81–83^. However, the complete mechanism by which the RPE participates in immunomodulation has not been fully elucidated.

Loss of PEDF is associated with aging and reductions in RPE functionality ^11,84^. Here, we accessed the potential immunomodulatory effects of the secreted homeostatic marker, PEDF, on damage outcomes and inflammation. Previous studies have described overexpression of or supplementation with PEDF as protective of photoreceptors and motor neurons, improvements in mitochondrial function and cortical neurons after damage, and inhibition of inflammatory damage ^2,22,45,85–88^. Additionally, deletion of PEDF is associated with aging, increased inflammation, and increased loss of visual function ^3,20,24,89^. Our results confirm the findings of other studies since the loss of PEDF resulted in increased retinal thinning, more damage-associated auto-fluorescent dots at the RPE-photoreceptor interface, significant loss of the photoreceptor layer, and increased cell death compared to littermate controls.

Additionally, when evaluating the retinal function, we found that the RPE of PEDF ^KO/KO^ animals had a reduced capacity for rhodopsin metabolism after LIRD compared to littermate controls at the same time point. Retinal function loss reduced scotopic *a*-,*b*, and *c*-wave amplitudes by five to seven days after light damage in PEDF ^KO/KO^ animals compared to littermate controls. These data suggest that PEDF is protective against excessive damage after phototoxic light exposure.

The RPE is the major contributor to IGF-1 secretion in the ocular environment ^90^. The importance of IGF-1 as a neurotropic factor and a regulator of immune cell function has been described in the eye and other tissue types under normal and pathological conditions, like cancer and ischemia ^53,55,56,91–93^. Additionally, decreases in IGF-1 expression have been correlated with aging, increased damage, and apoptosis in eye and brain studies^94–96^. To assess how the loss of PEDF may affect the expression and abundance of the neurotrophic factor, IGF-1, we first evaluated IGF-1 immunoreactivity in retinal sections of PEDF ^KO/KO^ animals compared to PEDF ^+/+^ animals at baseline. Baseline data showed no significant changes in IGF-1 expression between genotypes. However, after insult, there was a considerable loss in IGF-1 expression beginning on Day 3 of PEDF ^KO/KO^ animals, which increased to Day 7. We confirmed these findings via western blot analysis, showing a significant reduction in IGF1 protein expression in PEDF ^KO/KO^ compared to wild-type littermates at day 7. IGF-1 inhibits apoptosis of photoreceptors via the downregulation of caspase-3 and c-JUN signaling; thus, the reduced expression of IGF-1 may explain the increased degree of apoptosis observed in Fig.3B ^53,95,96^. The presence of IGF-1 and Galectin-3 co-expression in neuroprotective immune cells has been reported previously ^55,56,97^. We also assessed the presence of IGF-1 in recruited subretinal immune cells adhered to RPE flat mounts collected from PEDF^+/+^ and PEDF ^KO/KO^ animals at day seven post-LIRD. We found that PEDF ^KO/KO^ animals had fewer IGF-1 positive immune cells (See Fig. 6W-Y) compared to the PEDF ^+/+^(Fig.6T-V) at the same time point. IGF-1 modulates macrophage responsiveness and activity when challenged with a high-fat diet, shifting the transcriptional and morphological phenotypes to that of an M2-like proinflammatory macrophage^98^. A decrease in IGF-1 and PEDF expression has also been described in aging studies, which may suggest a similar mechanism as observed during our light damage experiments in the absence of PEDF ^99–102^. The loss of IGF-1 expression with age likely affects microglia function and sensitivity. The loss of PEDF leads to insult-initiated down-regulation of IGF-1 protein expression and reduced recruitment of IGF-1-expressing immune cells.

Multiple groups have described a unique subclass of immune cells enriched in galectin-3, recruited during neurodegeneration in the brain and the eye ^33,97,103–106^. In the eye, galectin-3 enriched subretinal immune cells are recruited to the photoreceptor-RPE interface, suggesting that there may be a functional requirement for galectin-3 in the subretinal space ^64,105,106^. Elevated galectin-3 expression is associated with poor prognostic outcomes^65–67,70^. Additionally, an ocular proteome study comparing AMD patients to age-matched controls found a significant increase in the secretion of galectin-3 binding protein and pigment epithelium-derived factor from the RPE ^107^. However, the correlation between PEDF expression, Galectin-3 levels, and damage outcomes has yet to be investigated. We hypothesized that loss of PEDF will increase galectin-3 expressing cells and global expression of galectin-3, ultimately leading to increased inflammation in the ocular microenvironment. To evaluate this, we quantified the number of galectin-3 expressing cells that adhered to the RPE at baseline and day seven between PEDF ^KO/KO^ compared to littermate controls. At baseline, there was no difference between genotypes. However, after damage, we found that the total number of galectin-3 positive cells was significantly increased in PEDF ^KO/KO^ animals compared to wildtype controls (See Fig. 7A-E), suggesting that without damage, there is no increased infiltration of immune cells. However, after damage initiation, PEDF ^KO/KO^ animals had significantly more galectin-3 expressing cells infiltrating the subretinal space compared to wild-type littermates at the same time. Damage to the subretinal space, neurodegeneration, and aging are associated with an increased activation of inflammation signaling and recruitment of immune cells ^108–114^. To investigate if PEDF ^KO/KO^ animals exhibit differential expression of galectin-3 and inflammasome mediator NLRP3, we used digital drop PCR. We found that galectin-3 mRNA expression in the RPE from PEDF ^KO/KO^ was significantly reduced compared to wild-type littermates. However, after damage, there is a significant increase in Lgals3 and NLRP3 expression at day 7 in PEDF ^KO/KO^ animals compared to the wildtype controls, which dampens the expression of these genes at the same time point. In agreement with the gene expression data, galectin-3 protein expression was significantly lower in PEDF ^KO/KO^ animals compared to PEDF^+/+^. On day 7, post-damage, PEDF +/+ animals reduced galectin-3 expression considerably compared to baseline expression; conversely, galectin-3 increased in the PEDF ^KO/KO^ animals at the same time point. Notably, while galectin-3 protein expression in PEDF ^KO/KO^ animals increased over baseline expression, there was significantly lower expression of galectin-3 protein at day seven compared to PEDF ^+/+^. These data may suggest that loss of PEDF affects the steady state of galectin-3 expression. Interestingly, when we pharmacologically inhibited galectin-3 activity in PEDF ^+/+^ animals during LIRD, we found it significantly decreased PEDF levels compared to LIRD-only controls (see Fig. 10), leading to poorer visual outcomes (data not shown). These data suggest that PEDF protects ocular function after LIRD via a novel galectin-3-mediated mechanism. PEDF suppresses eye diseases and cancer studies ^58,59,115^. In this study, we hypothesized that the protective role of PEDF in the ocular microenvironment after damage includes regulation of inflammation and immune privilege via galectin-3 mediated signaling. This study reports a putative relationship between galectin-3 and PEDF, suggesting that galectin-3 enriched immune cells within the subretinal space are a positive regulator of PEDF expression after light damage. However, the precise molecular signaling by which loss of PEDF impacts Galectin-3 and IGF-1 expression requires further study.

**Figure 10:**
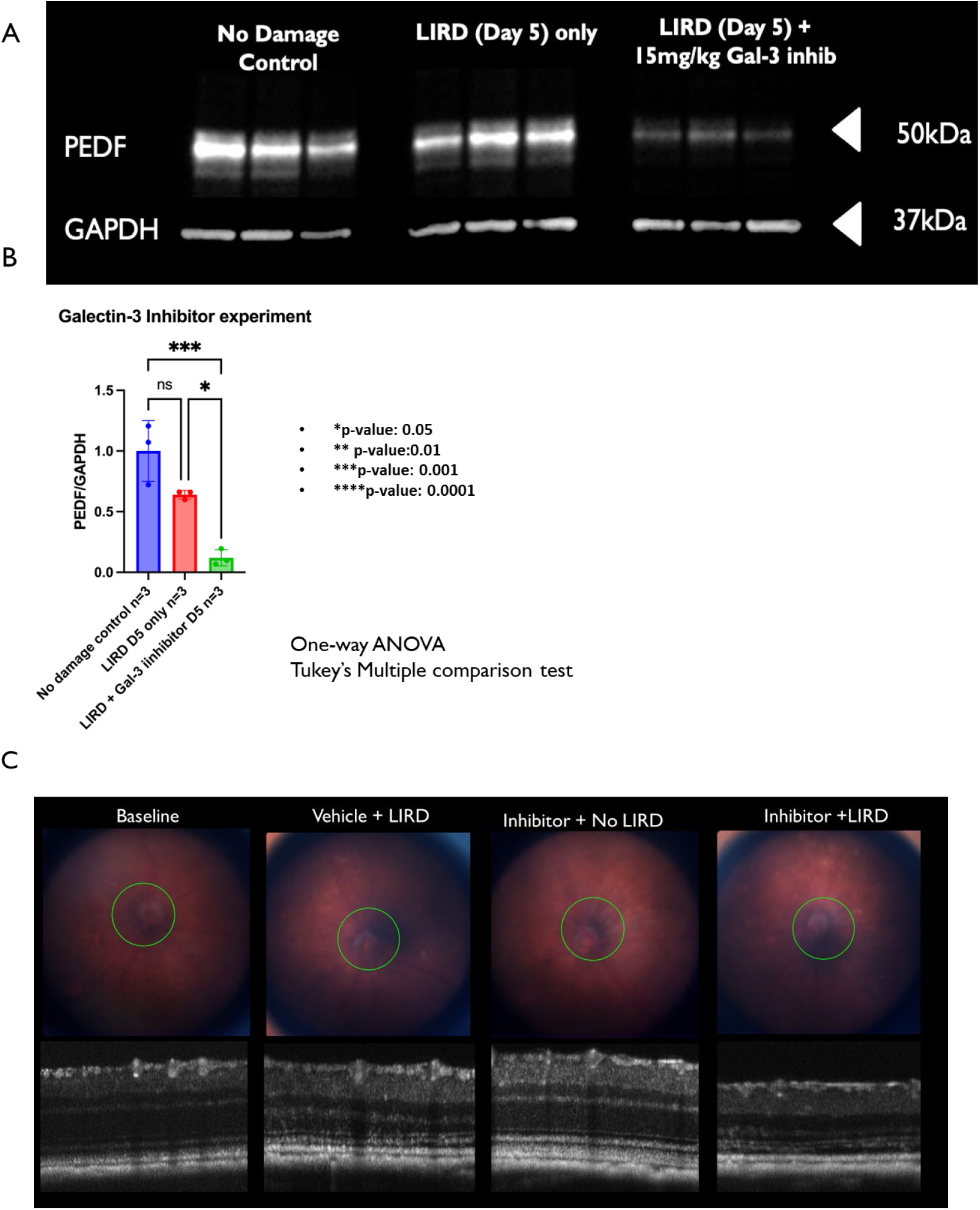
Treatment with Galectin-3 inhibitor, TD139, significantly decreases PEDF expression in PEDF ^+/+^ animals after LIRD. Figure 10A shows a western blot exhibiting that PEDF+/+ with no damage controls have high levels of PEDF, and exposing PEDF+/+ animals to LIRD shows a decrease in PEDF levels. Still, it is not significantly different from no-damage controls. However, by adding the galectin-3 inhibitor to LIRD, there is a significant loss of PEDF compared to LIRD, and there is no damage control. 10B is a quantification of 10A. (One-way ANOVA with Tukey’s multiple comparison test. No damage vs. LIRD only (Day 5 post) ns; not significant. No damage vs. LIRD (day 5) + Gal-3 inhibitor ***p-value<0.001. LIRD only (Day 5) vs. LIRD (Day 5) + Gal-3 inhibitor (*p-value<0.0.5). 10C shows representative fundus and retinal images taken using SD-OCT, displaying the effects of TD139 treatment with and without LIRD. Treatment with an inhibitor in conjunction with LIRD significantly increased retinal thinning compared to the control of LIRD only. Note: TD139 treatment alone does not affect visual function Galectin-3 expression levels ( Supplemental Figure 1).

**Figure 11:**
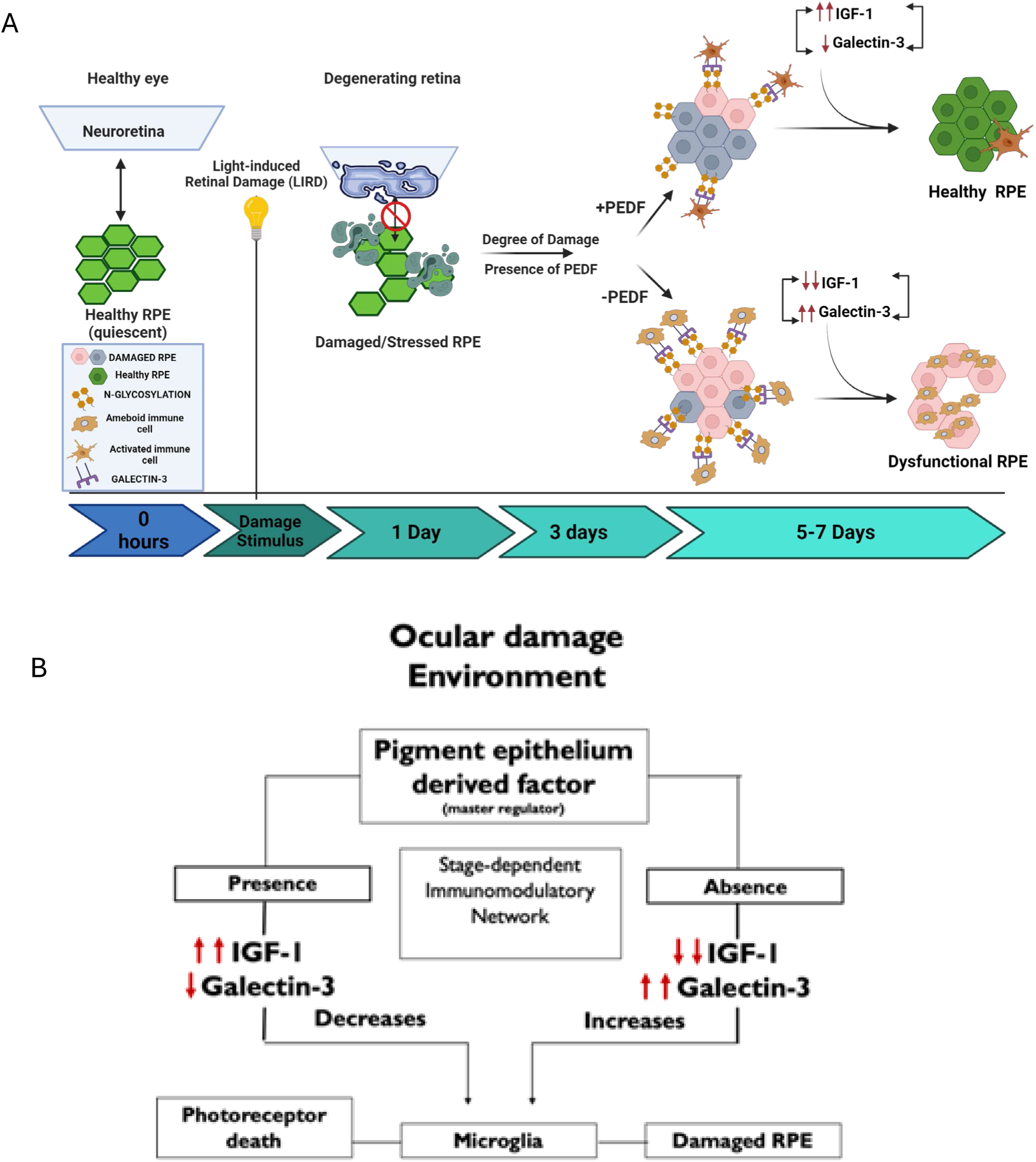
Schematic of Model Summary. Fig 11A: Schematic summary illustrating significant differences between PEDF ^+/+^ and PEDF ^KO/KO^ animals and the impacts on IGF-1 and Galectin-3 expression. Fig 11B: Shows the proposed immunomodulatory network influencing photoreceptor death, immune cells, and RPE cells. Images made using Biorender.

## Acknowledgments

Supported by Grants from the National Institutes of Health (R01EY028450, R01EY021592, P30EY006360, U01CA242936, R01EY028859, T32EY07092, T32GM008490); by the Abraham J. and Phyllis Katz Foundation; by grants from the U.S. Department of Veterans Affairs and Atlanta Veterans Administration Center for Excellence in Vision and Neurocognitive Rehabilitation (RR&D I01RX002806, I21RX001924; VA RR&D C9246C); and an unrestricted grant to the Department of Ophthalmology at Emory University from Research to Prevent Blindness, Inc. We would also like to thank Dr. Hans Grossniklaus and Dr. Sue Crawford at Northwestern University Feinberg School of Medicine for gifting the PEDF knockout mice used in this study.

