## Supplemental Figure 1 for "Loss of Pigment Epithelium Derived Factor Sensitizes C57BL/6J Mice to Light-Induced Retinal Damage"

5.7.1 **Supplemental Figure 1: Treatment with TD139 Alone does not Deleteriously Affect Visual Function or Galectin-3 Protein Levels**

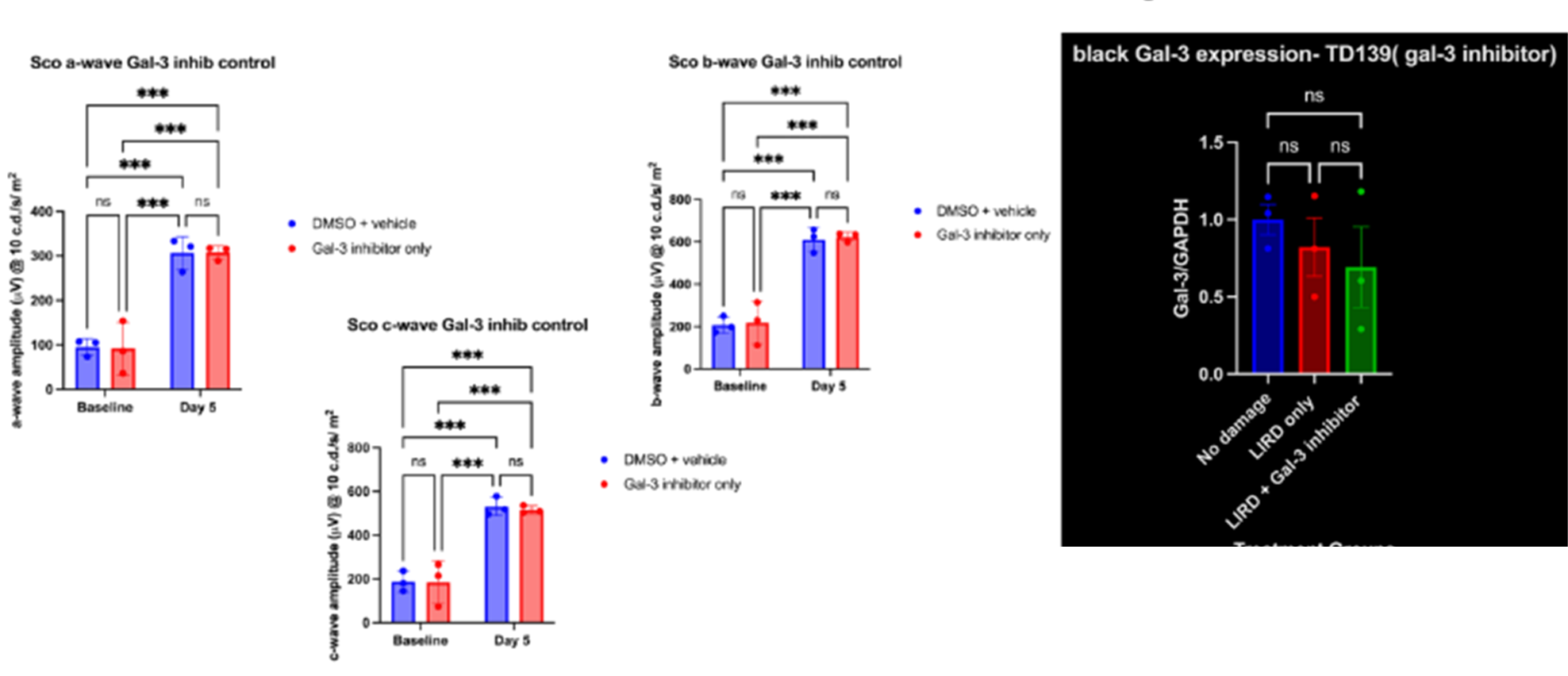

D

C

B

A

**Supplemental Figure 1: Treatment with TD139 Alone does not Deleteriously Affect Visual Function or Galectin-3 Protein Levels**

When comparing animals at baseline versus after 5 days of Galectin-3 inhibitor (TD139), we found that between the DMSO vehicle control and the experiment TD139 treated animals, there was not a statistically significant difference at baseline or at Day 5 (See figure S1 A-C). Additionally, when assessing total Gal-3 expression normalized to GAPDH of no damage, LIRD, only, and LIRD+ Gal-3 treated animals, the inhibitor did not deplete Galectin-3 levels, suggesting that its mode of function does not degrade Gal-3 protein. Taken together, these data show that inhibiting Galectin-3 activity using TD139 alone does not have deleterious off target effects on visual signaling or Gal-3 protein.
